# Three Essential Resources to Improve Differential Scanning Fluorimetry (DSF) Experiments

**DOI:** 10.1101/2020.03.22.002543

**Authors:** Taia Wu, Joshua Yu, Zachary Gale-Day, Amanda Woo, Arundhati Suresh, Michael Hornsby, Jason E. Gestwicki

## Abstract

Differential Scanning Fluorimetry (DSF) is a method that enables rapid determination of a protein’s apparent melting temperature (Tm_a_). Owing to its high throughput, DSF has found widespread application in fields ranging from structural biology to chemical screening. Yet DSF has developed two opposing reputations: one as an indispensable laboratory tool to probe protein stability, another as a frustrating platform that often fails. Here, we aim to reconcile these disparate reputations and help users perform more successful DSF experiments with three resources: an updated, interactive theoretical framework, practical tips, and online data analysis. We anticipate that these resources, made available online at DSFworld (https://gestwickilab.shinyapps.io/dsfworld/), will broaden the utility of DSF.

## Introduction

Differential Scanning Fluorimetry (DSF) is a biochemical assay used to measure the apparent melting temperature (Tm_a_) of a purified protein. In a typical DSF experiment, a protein solution is combined with a dye (*e.g.* SYPRO Orange) and heated in a qPCR instrument^1^. The DSF dye is minimally fluorescent in solution, but fluoresces brightly when bound to the hydrophobic regions of unfolded proteins. Therefore, as the protein unfolds during heating and reveals binding sites for the dye, fluorescence increases proportionally to unfolded protein abundance. The Tm_a_ is then calculated as the midpoint of the resulting fluorescence versus temperature curve, and higher Tm_a_s suggest a more stable native protein. Because Tm_a_s are broadly sensitive to changes in the biochemical state of the native protein, such as the binding of ligands^1–7^, mutations^8,9^, and proteinprotein interactions^10,11^, DSF is widely used to monitor diverse biochemical events via the associated change in Tm_a_ (ΔTm_a_) ^12,13^.

Compared to previous methods of Tm_a_ determination, DSF excels in accessibility and throughput. For example, circular dichroism (CD)^14^ and differential scanning calorimetry (DSC) ^15,16^, the biophysical gold-standard techniques for Tm_a_ determination, each require dedicated instruments, 50 μL to 2 mL of high-micromolar protein solution, and typically analyze a single solution at a time. In contrast, DSF requires little-to-no specialized instrumentation, 1 to 20 μL of low-micromolar protein solution, and accommodates throughputs from PCR tubes to 1536-well microtiter plates.

Given these advantages, DSF is used widely by labs in academia, government, and industry alike. For example, DSF may rapidly identify stabilizing buffer condition,, enabling structural characterizations^17–19^ or formulations of biotherapeutics^20^. High-throughput DSF screens are used to identify candidate binders to proteins of interest independent of enzymatic activity^7,12,21^. DSF has also been used to search for small-molecule correctors for genetically destabilized proteins^22–25^ which underlie many misfolding diseases^13,26,27^ such as cystic fibrosis^28,29^, Gaucher’s disease^30^ and Fabry’s disease^31^. For example, our lab conducted a primary chemical screen by DSF to identify ligands for cataract-associated forms of alpha B-crystallin^32^. These applications and others have been excellently reviewed^12,33^.

Yet DSF regularly fails. Sometimes a validated ligand induces no ΔTm_a_, a replicated experiment produces different results, the data cannot be fit to a simple sigmoid, and so on. While excellent resources exist guiding applications of successful DSF, as well as some descriptions of its systematic shortcomings^33,34^, there is currently little to no framework for optimizing failed DSF conditions into successful ones, let alone how to do so efficiently. Instead, failures and solutions are passed on at conferences or group meetings if at all, so when an experiment fails, it is often unclear what to try next.

The purpose of this paper is therefore to guide readers from failed to successful DSF. This begins with re-joining DSF with its theoretical underpinnings—a combination of unfolding thermodynamics and kinetics, and dye-binding—in a convenient and usable manner. In the first of three resources, we start with an empirical assessment of the ability of the current theoretical framework to describe real DSF data, and find that it overlooks two widespread features which carry important practical ramifications: kinetic influence on protein unfolding, and atypical dye activation. We present a correspondingly-updated theoretical framework and associated computational model for DSF, and demonstrate its increased ability to describe widely-observed empirical phenomena in DSF at odds with the current framework. We make this model available as an interactive online tool at DSFworld (https://gestwickilab.shinyapps.io/dsfworld/). In the second resource, we build on this updated framework to identify common but largely undescribed technical pitfalls which ruin even well-designed DSF experiments, and present experimental bestpractices to minimize them. Finally, we provide a free DSF data analysis software at www.DSFworld.com. DSFworld addresses major existing limitations in data, including customizable visualizations based on user-defined experimental variables, as well as streamlined handling of complex, multi-transition data. Together, we hope that these three resources—theory, practical tips and data analysis—will help the community perform more successful DSF experiments.

## Results

### Section I: An improved theoretical framework for DSF

In a DSF experiment, the measured fluorescence signal is a product of multiple molecular events, including protein unfolding, dye binding, and dye activation (*e.g.* an increase in quantum yield). The current theoretical framework (termed here “Model 1”) underlying the design and interpretation of DSF experiments includes two major assumptions: (i) that protein thermal unfolding reflects thermodynamic equilibrium and (ii) that dye fluorescence provides a proxy for unfolded protein abundance.

Here, we demonstrate that this widespread theoretical framework for DSF falls short in two straightforward, yet important, ways. First, we show experimental and theoretical evidence that the kinetics of protein unfolding, not just the thermodynamics, influences the outcome of most DSF experiments. Second, we show that dye binding is not always exclusive to the unfolded state; rather, the dye sometimes binds to the native state. In other cases, the dye fails to bind the unfolded states, such that it cannot accurately reveal the unfolding process. Incorporating these considerations, we propose an improved theoretical framework for DSF, termed “Model 2”.

Currently, interpretation of most DSF experiments assumes that protein unfolding is dominated by thermodynamic contributions, yet the results of extensive CD and DSC studies have suggested that kinetics plays a major role, especially for thermal unfolding^35,36^. Based on this classical literature, we hypothesized that DSF results might also include a contribution from kinetics. If so, calculated Tm_a_ values measured by DSF would be expected to depend on the rate at which the sample is heated (°C / min). To test this idea, a model thermodynamic unfolder (hen egg white lysozyme) ^37^ and a model kinetic unfolder (malate dehydrogenase; MDH)^38^ were analyzed by DSF at systematically increased heating rates (0.5, 1, 2 and 4 °C/min; Figure 1a). As expected, we found that the calculated Tm_a_ value for lysozyme (69 °C) was largely unaffected by heating rate (ΔTm_a_ of only 0.8 °C, calculated between the extreme heating rates of 0.25 and 2 °C/min; Figure 1b, c). Conversely, a strong effect was observed for MDH (ΔTm_a_ = 3.1 °C; Figure 1b, c). To understand how widespread this kinetic contribution might be, we tested six additional proteins that vary in structure and molecular mass (PPIF, PerAB, PPIE, Per2, Hsc70, CHIP/STUB1; Supplemental Table 1). Strikingly, the Tm_a_ values for all six proteins, like MDH, also varied substantially with heating rate (ΔTm_a_ = 2.2 to 5.4 °C; Figure 1b, Supplemental Table 1). This result suggests that kinetics does significantly influence the outcomes of many DSF experiments.

**Figure 1.**
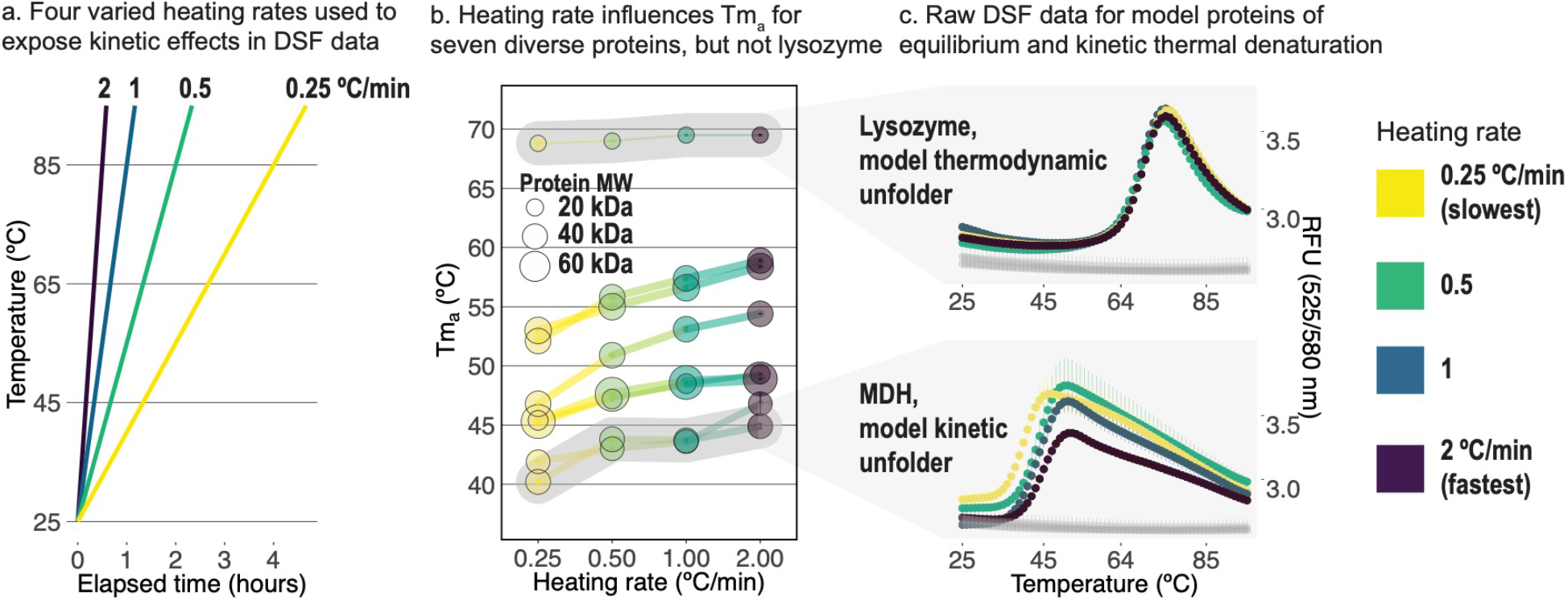
The kinetics of protein unfolding impact DSF results. a. Schematic of the four thermocycling protocols used to generate results presented in b, highlighting the different elapsed times required to reach the same temperature between them. b. From the diverse eight protein panel, seven display heating-rate dependent changes in Tm_a_, similar to the model irreversible unfolder malate dehydrogenase (MDH). Only the model reversible unfolder lysozyme shows heating rate-independent Tm_a_s. c. Raw DSF data for model equilibrium unfolding protein lysozyme and model kinetic unfolding protein MDH at the four ramp rates. No-protein controls (5X SYPRO Orange in buffer) are shown in grey. Results presented are the average of from three technical replicates +/-standard deviation.

To incorporate unfolding kinetics into the theoretical framework of DSF, we chose the simplest of the classic Lumry-Eyring models which combine thermodynamics and kinetics of unfolding: ^39^

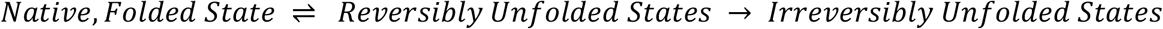

The use of this model is informed by pioneering work on the analysis of DSC experiments^35^. However, we had to adapt it for use in simulating DSF data by including the contribution of dye binding. Specifically, we multiplied the abundance of each protein state F(T) and the kinetic partitioning between these states, L(T), by the extent to which they each activate the dye D(T). Importantly, the D(T) term is a product of both a dye’s binding affinity and its quantum yield. Thus, in Model 2, the measured fluorescence is predicted to be a sum of the RFU contributions from dye binding to each of the three states (RFU_native_, RFU_reversibly unfolded_, RFU_irreversibly unfolded_), and corrected for the empirically observed temperature-dependent losses in fluorescence, yielding the final model:

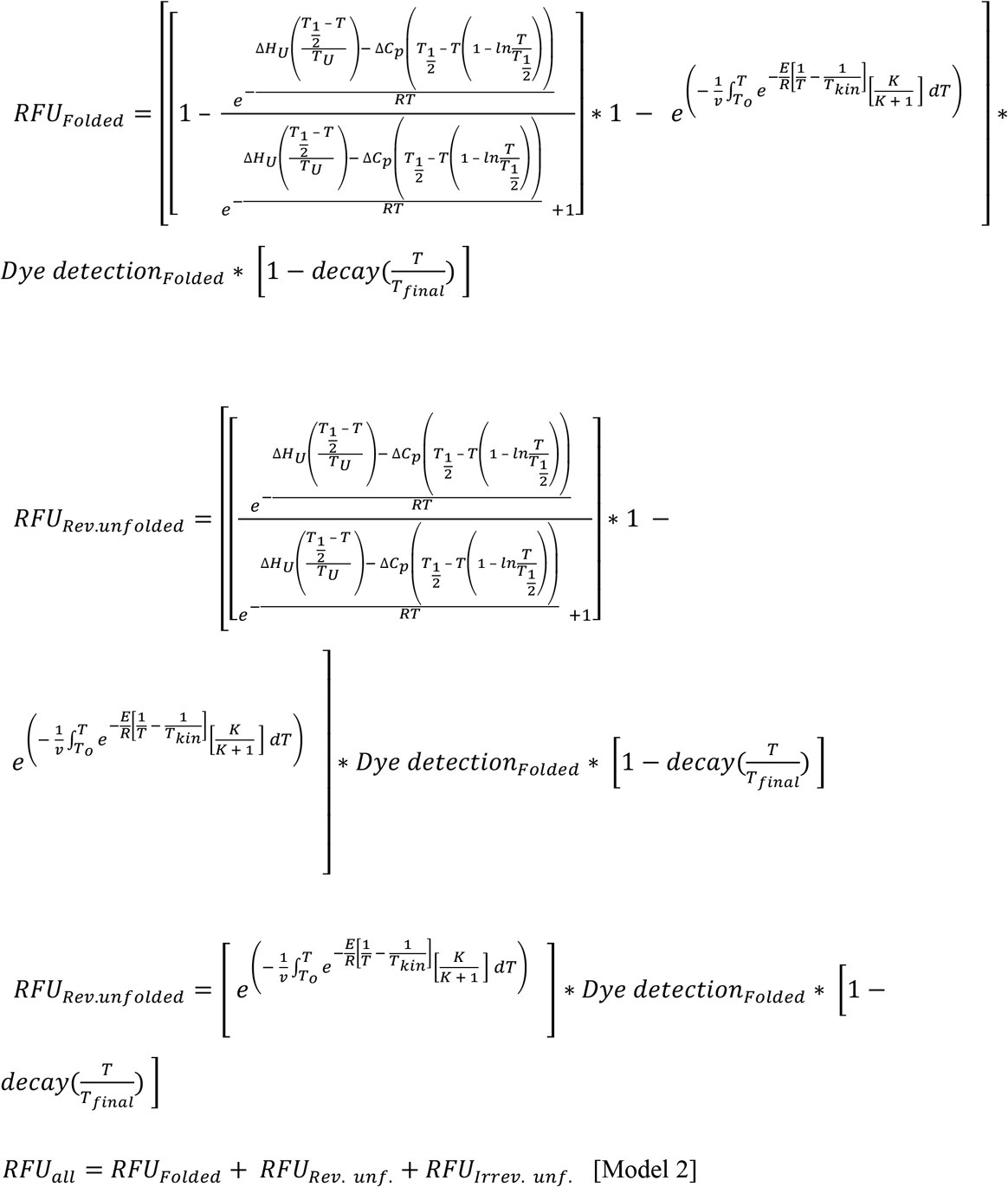

As a head-to-head test, we simulated DSF results at varied heating rates from either Model 1 (the current DSF theoretical framework) or Model 2 and compared them to the experimental data from Figure 1. As expected, we found that Model 1 could not account for the dependence of Tm_a_ on heating rate, while Model 2 successfully recapitulated it in magnitude and direction for both MDH (ΔTm_a_ = +2.2 to 5.4 °C empirical; +6 °C theoretical), and the thermodynamic unfolder lysozyme (ΔTm_a_ = +0.8 °C empirical; +0.8 °C theoretical) (Supplementary Figure 1). This agreement supports the utility of Model 2 for improved interpretation of DSF data.

We next tested whether Model 2 could recapitulate two additional, widespread conundrums in DSF experiments. First, we explored the issue of validated ligands producing negative ΔTm_a_ values. In the thermodynamic Model 1 framework, selective ligand binding to the native protein must increase Tm_a_, such that calculated ΔTm_a_ values should always be positive. However, negative ΔTm_a_ values appear routinely in published reports, even when the ligand is properly validated by other techniques^21,32,40–42^. How is this possible? Using Model 2, we generated a theoretical DSF dataset, in which we varied the impact of ligand binding on the activation energy of protein unfolding (Ea), and found that decreased Ea values were sufficient to increase kinetic partitioning into an irreversible unfolded state and produce negative ΔTm_a_ values (Figure 2c). Thus, Model 2 was able to produce theoretically sound reason for why negative ΔTm_a_ values are sometimes observed. We also noted that these negative ΔTm_a_ values were accompanied by a systematic decrease in the slope of the transition (Figure 2d), suggesting that analysis of curve shape, not just ΔTm_a_, is both theoretically-justified and potentially advantageous. At the least, these results prompt re-evaluation of the current practice of discarding all negative ΔTm_a_ hits from high throughput DSF screens.

**Figure 2.**
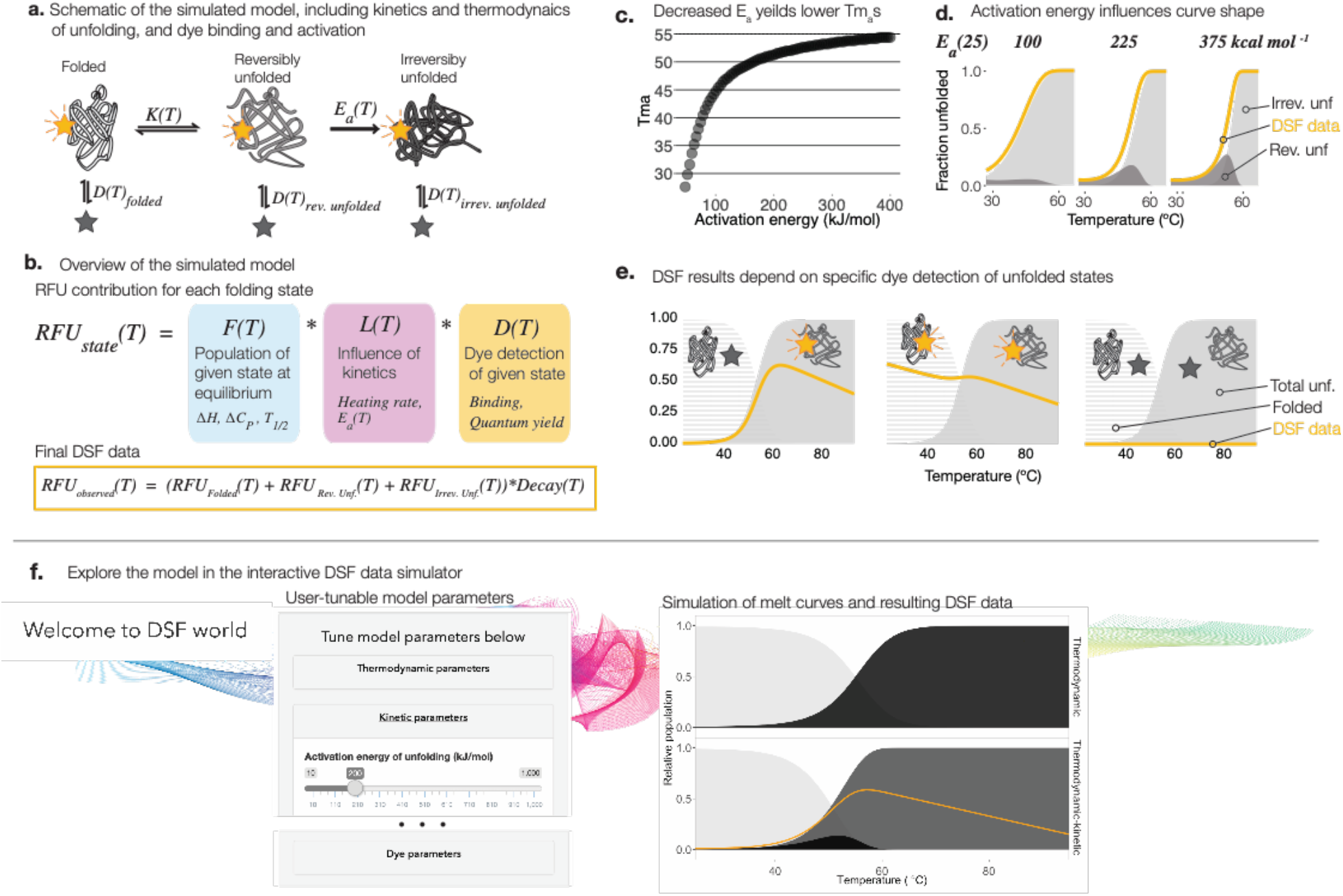
An updated thermodynamic-kinetic DSF data model “Model 2” recapitulates empirical observations and motivates a nuanced interpretation of DSF results. a. The mixed thermodynamic-kinetic unfolding model used, comprising an initial equilibrium “thermodynamic” step followed by an irreversible “kinetic” step. Dye detection is quantified for each folding state individually. b. Schematic of model construction. Top: The contribution of each folding state to the final simulated DSF data. The relative population of a state is given by its equilibrium population K(T), modified by kinetic influence L(T). RFU is the product of the population of a state and the extent to dye detects it D(T). Bottom: The observed DSF data is obtained by summing RFU contributions for all states followed by multiplication by a protein-independent Temperature-dependent decay in fluorescence. c. Decreased activation energy of unfolding is sufficient to decrease Tm_a_. d. DSF curve shape is significantly altered by changing only activation energy of unfolding (Ea(T)). e. Model simulations of different DSF data obtained for the same unfolding trajectory as the dye detection of the various states changes. Left: ideally, dye detects only unfolded states. middle: dye detection of folded states gives high initial RFU and an obscured transition. right: no detection of unfolded states produces no transition. f The model presented in this paper can be explored further in the Interactive Modeling section of DSFworld https://gestwickilab.shinyapps.io/dsfworld/.

Next, we examined why, for some proteins, DSF fails to reproduce the melting curves that are expected from CD or DSC. For example, it is relatively common to encounter DSF curves in which the fluorescence starts high at low temperature and then decreases, rather than increases, during heating. We hypothesized that this effect might be due, in part, to aberrant binding of the dye to the native, folded state. To test this idea, we generated a theoretical DSF dataset from Model 2, in which the affinity of the dye for protein states—native, reversibly unfolded, and irreversibly unfolded—was individually enhanced. We found that, when dye was activated by the native state, the resulting Tm_a_ values were aberrantly elevated (Figure 1 c), matching what is often observed in practice. In the other extreme, when dye was not activated by the unfolded states, the fluorescence was “flat” and unfolding transitions were not detectable (Figure 2e, right panel). These results also agree with our empirical observations of some proteins (Supplemental Figure 3). Extending this concept to data analysis, we found that including a temperature-dependent initial fluorescence population in the fitted sigmoid model improves the accuracy of Tm_a_ values calculated for proteins with native-state dye activation (described in more detail in Section III). Finally, extending this concept to the bench, we found that elimination of extraneous sources of dye activation is a critical step in optimizing DSF conditions (described in more detail in Section II).

An interactive, online tool for comparison of Models 1 and 2 is available at DSFworld and the associated R script is publicly available on GitHub.

### II. Practical tips, theoretically grounded

Although the theoretical framework described in Section I appears to be a useful tool to improve DSF experimental design and interpretation, it does not account for experimental artifacts. A separate consideration of artifacts is important because common ones can produce effects on fluorescence that qualitatively resemble those introduced by using Model 1. In this section, we describe potential sources of these artifacts, alongside five best-practices to avoid them. In addition, to reducing the impact of these artifacts, the other goal of this section is to improve reproducibility and maximize sensitivity of DSF experiments.

#### i. Include no-protein controls for every condition

Because DSF relies on the use of dye fluorescence as a proxy for protein unfolding, identifying and minimizing sources of protein-independent fluorescence is a critical step. Indeed, the fluorescence of SYPRO Orange is known to be sensitive to excipients that are common in biological buffers, such as glycerol, detergents, lipids and EDTA^34,43^. Here, we focus on two especially pernicious and common sources of protein-independent fluorescence: dye binding to colloidal aggregates and dye binding to the plastic used in manufacture of some microtiter plates.

Dye binding to plastic is a common problem in DSF. This artifact manifests as significant fluorescence in the absence of any protein (Figure 3a). In extreme cases, this artifact produces a fluorescence transition that, upon first inspection, mimics the shape of a curve that might result from dye binding to a folded protein (Figure 3b). One can readily discriminate between these possibilities by testing dye fluorescence in buffer without protein. An under-appreciated aspect of this artifact is that it varies between plate lots (*e.g*. microtiter plates manufactured at different times). Thus, because offending plates might have the same catalog number and vendor, each new lot should be tested for protein-independent dye-activation before use (see Supplemental Figure 4 for a plate compatibility test protocol).

**Figure 3.**
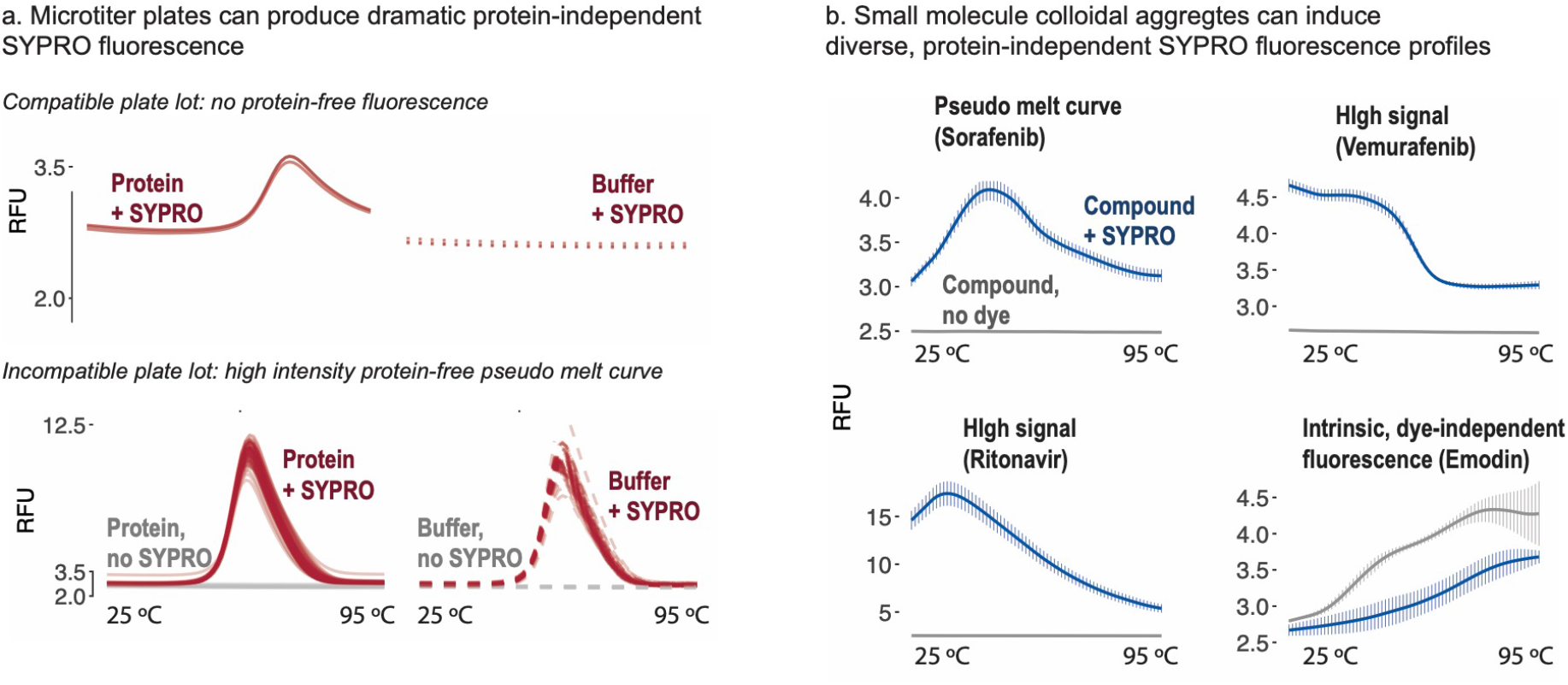
No protein controls reveal misleading protein-independent fluorescence. a. Spaghetti plots of DSF data from one DSF-compatible microtiter plate lot (Lot 1, left), and two DSF-incompatible plate lots (Lots 3, 3). In a DSF-compatible plate, 5X SYPRO produces no significant fluorescence at any temperature in the absence of protein. In incompatible plate lot 2 (left), 5X SYPRO in buffer alone produces “phantom” melting curves in all wells (red). In incompatible plate lot 3 (right), 5X SYPRO in buffer alone produces high, decaying signal in all wells (red). In both cases, protein-independent fluorescence is more than sufficient to distort or dwarf typical DSF data for the model protein lysozyme in a compatible plate lot (blue, left and right).

When a DSF experiment involves addition of a small molecule, we found that an additional control must be performed to reduce artifacts associated with dye activation by to the compound. Much like plate-related artifacts, these ones manifest as a protein-independent increase in dye fluorescence. We suspected that dye binding to colloidal aggregates might underlie some of these cases. This hypothesis is based on work describing the formation of colloidal aggregates by small molecules as a recognized mechanism of pan-assay interference compounds (PAINS)^44^–^46^. To test this idea, we assembled a panel of eight compounds that have been reported to form colloidal aggregates, but that otherwise vary in chemical structure (Supplementary Figure 5)^47,48^. Interestingly, seven of the eight compounds induced protein-independent fluorescence in the DSF experiment, which was sufficient to obscure the melting transition of lysozyme (Figure 4, Supplementary Figures 6 & 7). Importantly, we also found that the addition of 5X SYPRO Orange (equivalent to 10 μM, Supplementary Figures 8 & 9) further aggravated colloidal aggregation of all compounds (Supplementary Figures 6). Thus, dyes and compounds can sometimes combine to create significant artifacts. Together, these results implicate aggregates as another source of protein-independent dye activation.

**Figure 4.**
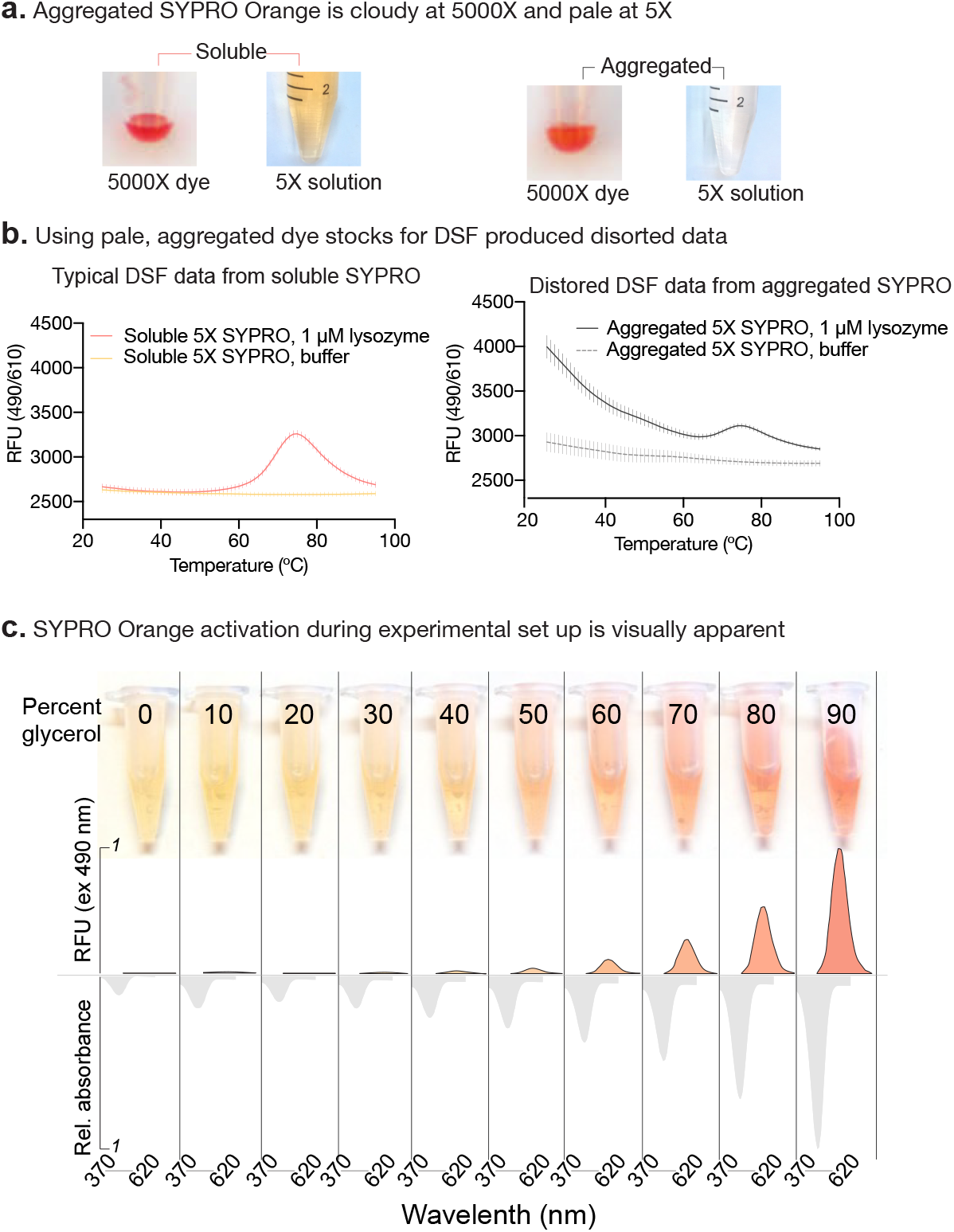
Atypical color suggests non-optimal conditions. a. Typical, soluble 5000X SYPRO Orange stock appears dark and clear, and produces orange 5X working solutions (left), while spontaneously aggregated freshly-thawed 5000X SYPRO Orange appears cloudy and light, and produces markedly pale 5X working solutions. b. DSF data collected for the model protein lysozyme using soluble 5X SYPRO Orange shows a clear, accurate melting curve and no protein-independent fluorescence (left). The same experiment performed using the pale, aggregated stock produces high room-temperature fluorescence and an obscured melting curve and increased protein-independent fluorescence (right). c. Room-temperature activation of SYPRO Orange, in this figure by increasing concentrations of glycerol, often produces pink-pigmented solutions during experimental set-up.

Beyond conducting “no-protein” and “compound-only” controls, what other steps should be taken to reduce the impact of artifacts? If possible, minimize the use of glycerol, which can produce viscosity-related changes in dye fluorescence. Likewise, the use of detergents, EDTA, or other additives should be minimized and any stock solutions fresh filtered before use. As a benchmark, we often use fresh solutions of 0.001% Triton X-100 or 1 mM EDTA without issue. It is also recommended that each experiment includes a benchmarked positive control. For example, we typically include a DSF standard composed of 10 μM hen egg white lysozyme and

10 μM (“5X”) SYPRO Orange in buffer (e.g. 10 mM HEPES, 200 mM NaCl pH 7.2; Tm_a_ ~ 69 °C). If this standard produces atypical data (*e.g*. high initial fluorescence, aberrant Tm_a_ value), this suggests a reagent-based artifact. Even with these precautions, it is sometimes difficult to completely eliminate all contributions of artifacts. For example, a protein might not be stable in the absence of detergent. In our experience, it is reasonable to proceed if the contribution of the artifact is less than 10% of the desired signal.

#### ii. Optimize heating rates

One practical implication of Model 2 is that both Tm_a_ values and curve shape are sensitive to the heating rate employed. In addition to the theoretical implications, this relationship is also practically useful: in our hands, both the reproducibility and sensitivity of DSF data can be optimized by varying the heating rate employed. For example, when a known ligand interaction produces small ΔTm_a_, the value can often be increased by heating at 20, 40, 60, or 120 seconds per degree. These adjustments balance the relative contributions of variables in Model 2, such as the kinetic component of unfolding, and allow enhancement in signal to noise ratio. Similarly, ligand-induced ΔTm_a_ values are often enhanced by increasing the ligand concentrations by 2 to 10-fold above the K_d_. This adjustment can compensate for temperature-dependent losses in binding affinity, which are often not included in common Kd measurements by isothermal calorimetry (ITC), surface plasmon resonance (SPR) or other methods. In the literature, reports of changes from 2 – 12 °C currently dominate, from 1 - 2 °C are also common. Changes exceeding 15 °C are rare for non-covalent interactions, though exceptions are possible--the binding of biotin increases the Tm_a_ of streptavidin by a remarkable 47 °C^5^. An important corollary of these observations is that Tm_a_ (or ΔTm_a_) values collected using different thermocycling protocols are not comparable. Not all commercial qPCR instruments cool efficiently or keep time accurately, so compared experiments should be performed on the same instrument when possible.

In addition to improving signal, optimized thermocycling protocols can also make DSF results more reproducible and easier to analyze. For example, shallow or poorly resolved transitions are difficult to fit using common methods, such as first derivate. When these data features are encountered, it is sometimes helpful to switch to up-down heating mode. In up-down mode thermocycling, the reaction is re-cooled to 25 °C between heating increments (Supplemental Figure 2), and fluorescence is measured at the low-temperature steps. As implied by Model 2, up-down heating monitors exclusively irreversibly unfolded states, providing a potentially simplified portrait of otherwise complex unfolding transitions.

#### iii. Optimize buffers and additives to stabilize the protein

A common problem in DSF experiments is the presence of protein aggregates in the starting sample. Often, this issue appears as high initial fluorescence, likely due to binding and activation of dye by the contaminating, aggregate. In this case, filtering typically removes the aggregate and the interfering signal. However, if the problem persists, dye may be detecting the folded protein (as discussed in Section I, Figure 2e). To minimize the contribution of this event, buffer conditions can sometimes be varied to stabilize the protein and/or occlude the dye binding sites. For example, adding co-factors (*e.g*. ADP), coordinating metals (*e.g*. LiCl, MnCl2) ^49^, or endogenous ligands can sometimes help, perhaps by competing for dye-binding sites. Another approach is to stabilize the protein by adjusting the ionic strength of the buffer and/or including additives, such as DMSO or sucrose.

#### iv. Heed visual cues

During experimental set-up, trust your eyes. DSF dyes absorb in the visible spectrum, and while minor variations in visual pigmentation and intensity are normal, significant deviations often portend unreliable results. For example, aggregation of the SYPRO Orange in stock solutions can occur regardless of storage conditions (*i.e*. anywhere between 25 °C and −80 °C). Aggregated SYPRO Orange appears as a solution with dull pigmentation (Figure 4a), and, if used, it produces both protein-independent fluorescence and muted transitions (Figure 4b, right). Fresh stocks should be made and filtered to remove aggregates. Similarly, look for changes in color for any solution, especially when a new buffer, additive or small molecule is introduced. SYPRO Orange loses both fluorescence in DSF as well as room-temperature pigmentation below ~pH 5 (Supplementary Figure 10). Conversely, protein-independent SYPRO Orange fluorescence induced by small molecule additives often corresponds to an increase in visual pigmentation (Supplementary Figure 10).

#### v. Know when to change assays

DSF isn’t right for all questions, or all proteins. For example, ITC^50^, SPR^51^ or similar platforms are likely to be superior methods to determine the K_d_ of ligand binding. Similarly, chemical (rather than thermal) denaturation is a better way to determine the ΔG_unfold_ for a protein, because it provides a closer approximation of equilibrium unfolding. Rather, DSF excels in accessibility and throughput. Thus, if other biophysical methods do not provide a K_d_ or ΔG_unfold_ for the system of interest, than it is unlikely that DSF will “fix” the issue. Rather, any apparent change in Tm_a_ could be the result of one of the artifacts mentioned above.

### Section III. Data analysis

Inaccessibility of robust, efficient data analysis remains a significant and widespread bottleneck for DSF users. Recent reports have presented both scripts and websites for the analysis of DSF data^52–57^, we found that two substantial bottlenecks remained unaddressed. First, there remains a need for tools to efficiently, and flexibly visualize the effects of experimental variables to facilitate data interpretation and optimizations. At DSFworld, we address this issue in the following manner (Figure 5): raw RFU data can first be uploaded either exactly as exported by most standard qPCR instruments, or as a csv file with temperature in the left-most column and data in the columns to the right. Then, any number of experimental variables (*i.e*. ligand, ligand concentration, pH, and protein) can be efficiently assigned using a provided plate-layout template. The user-defined experimental variables become automatically available for use in the generation of customized plots. Wells which are identical in all assigned variables are automatically considered replicates and can be plotted as averages with standard deviations. Together, these features facilitate rapid, flexible comparison of multiple experimental variables. Plot aesthetics such as text size and titles can be customized, and final plots can be downloaded in high resolution as pdfs.

**Figure 5.**
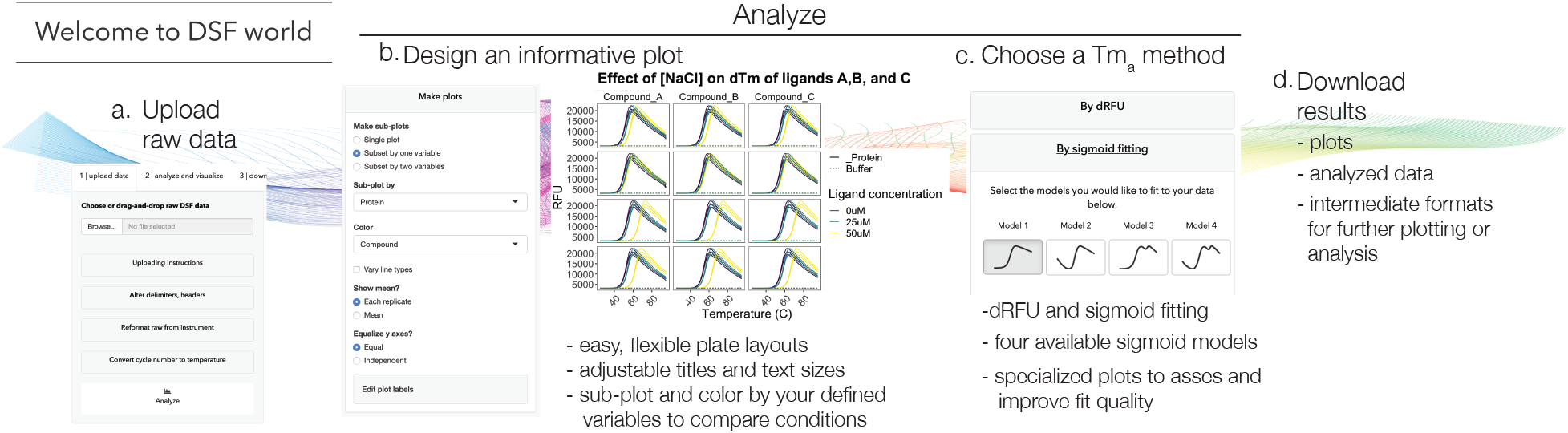
Online data visualization and analysis at DSFworld. a. Upload raw DSF data exactly as exported for supported instruments, or with Temperature in the left-most column and RFU data to the right. b. If desired, define replicates and experimental conditions by editing the names table directly, or uploading a custom csv plate layout (see template) to define any number of variables (e.g. compounds, concentrations, buffers). Rapidly visualize the effect of each condition tested in the “make plot” panel by sub-plotting and/or varying colors and line types by any user-defined variables. In this mock example, the effects of NaCl concentration on Tm_a_ sensitivity to the binding of ligands is explored. Set custom titles and text size for publication- and presentation-quality figures. c. Choose a Tm_a_ method, either dRFU or one of four sigmoid fits. c. Download Tm_a_s, plots, and various formats of raw and analyzed data for publications, presentations, or further analysis.

The second bottleneck we attempt to address at DSFworld is the availability of flexible and robust curve fitting, particularly for complex or multi-component transitions. To create an analysis pipeline rooted in both the theoretical framework of DSF and the diversity present in real DSF data, we began by assembling a panel of 35 purified proteins of diverse size, structure, and oligomeric state, and then performed DSF experiments on each one in multiple conditions (*e.g*. buffer, ligand) to generate 347 individual DSF curves (Supplementary Figure 11). Drawing from the implication from Model 2 that DSF data analyses should allow for both room-temperature dye binding, we composed four sigmoid models for fitting: a single or double decaying sigmoid (fits 1 and 3), or a single or double decaying sigmoid with high initial fluorescence (fits 2 and 4). Strikingly, we found that 341/347 curves were well described by at least one of the four fits (see Supplementary Note 2). At DSFworld, Tm_a_ values can be determined by either the maximum of the first derivative, or any of the four fits. After fitting, the best model can be either selected manually from an interactive plot, or automatically based on the Bayesian Information Criterion. The full analysis workflow, from raw data uploading to results downloading, is available at DSFworld^58^–^68^ and as stand-alone scripts and modular web applications on GitHub.

## Discussion

Here, we present three resources to help users design, optimize and troubleshoot DSF experiments: interactive theoretical modeling, practical tips and online data analysis. These efforts culminate in an on-line resource: DSFworld (https://gestwickilab.shinyapps.io/dsfworld/). In the first Section, we found that linking DSF experiments to established protein unfolding theory improved our ability to predict common problems and design potential solutions. For example, using a panel of seven proteins of diverse size, we observed that kinetics plays a significant role in the outcome of DSF experiments, motivating reconsideration of the thermodynamic framework that is widely used to-date. Accordingly, we present an updated theoretical model, Model 2, which includes attention to both thermodynamics and kinetics of unfolding. Using simulated results, we demonstrate that changes in the activation energy of unfolding (E_a_) effect curve shape and Tm_a_, providing a possible explanation for how legitimate ligands can sometimes decrease, rather than increase, ΔTm_a_. This is an important advance because such reports in the literature are often called into question as potential artifacts. Given the current interest in protein stability, a fresh, theory-driven approach to this question seems warranted.

These improved models illustrate that curve shape, not just Tm_a_, is a useful feature of DSF experiments. For example, ligand binding might be evident by a change of curve shape, even if the ΔTm_a_ is modestly affected. However, it is important to note here that the physical mechanisms responsible for this observation are often not clear. Going forward, it will be important to establish, using structural and computational methods, if there are specific elements of curve shape (*e.g*. slope, initial fluorescence, number of transitions) that are most informative. In the meantime, we suggest a broader view of DSF results than just a singular focus on Tm_a_.

Other fields of biophysical measurement, such as SPR, have benefitted from application of user-initiated, quality control criteria. These efforts often coalesce around shared, online resources. Towards that goal, we report an online DSF data visualization and analysis at DSFworld. As part of this effort, we include customizable data fitting, visualization and plotting features, which includes Tm_a_ calculation by first derivative or any of four sigmoidal models. Furthermore, we have made the full code for DSFworld available on GitHub, alongside both stand-alone scripts and modularized web applications for each of the individual data analysis problems resolved at DSFworld. This repository can serve as both a venue and resource for the continued improvement of challenges in DSF data analysis.

We hope readers can then use these resources—theory, technical tips, and data analysis— as a foundation to drive DSF forward through their own innovations, designing powerful experiments and completing them easily.

## Acknowledgements

The authors are grateful to Dr. Matthew Jacobson (UCSF) for productive conversations on the theoretical models presented, and Dr. James Fraser for constructive criticism. We thank Dr. Daniel Elnatan for productive conversations on and examples of DSF data fitting strategies. We are grateful to many members of the UCSF community for the beta-testing of DSFworld, especially Dr. Ziyang Zhang and Douglas Wassarman, and to Hadley Wickham for making its creation possible. This work was supported by an NSF fellowship to TW and a grant from the Tau Consortium to JEG.

## Author Contributions

Project design, manuscript preparation (T.W., J.E.G.), conduct experiments (T.W., J.Y., Z.G.D.), data interpretation (T.W., J.Y., Z.G.D., J.E.G.), experimental support (A.W., A.S., M.H.), laboratory management (J.E.G.).

## Materials and methods

For all procedures, no unexpected or unusually high safety hazards were encountered.

Differential Scanning Fluorimetry. Unless otherwise noted, conditions for all DSF experiments were: 10 μM protein; 10 μM (“5X”) SYPRO Orange, (Thermo Fisher Scientific Ref S6650, Lot 2008138) in 10 mM HEPES 200 mM NaCl pH 7.20 0.22 μm filtered (Millex-GS 0.22 μm sterile filter unit, ref SLGS033ss, lot R8EA61590), with a final DMSO concentration of 0.1% and final reaction volume 10 μL per well in a 384-well microtiter plate (Axygen PCR-284-LC480WNFBC Lot 09819000). DSF experiments were monitored in a BioRad CFX 384 qPCR in the FRET channel.

Dynamic Light Scattering. From 5 mM stocks in DMSO (Sigma Aldrich prod 276855-100 mL lot SHBK3913), six three-fold serial dilutions of each compound were prepared in DMSO (150 – 0.2 μM), diluted in 10 mM HEPES 200 mM NaCl pH 7.20, and combined with either buffer or lysozyme, with or without SYPRO Orange, to final concentrations of 10 μM lysozyme, 150 – 0.20 μM compound, 10 μM SYPRO Orange, 2.5% DMSO. All reagents were filtered prior to use Millex-GS 0.22 μm sterile filter unit, ref SLGS033ss, lot R8EA61590. 20 μL of each final solution was added to a DLS plate with no plate seal, and colloidal aggregation was measured using a Wyatt Technologies DynaPro Plate Reader II (acquisition time 2 sec, 20 acquisitions, with auto-attenuation, with temperature control, 25 °C). For the matched DSF experiments, 10 μL of the same solutions was added to each well of a DSF plate (Axygen PCR-284-LC480WNFBC Lot 09819000) and heated from 25 – 95 °C at a heating rate of 1 °C/min and monitored in the FRET channel of a BioRad CFX384. Experiments were performed in technical triplicate and presented as mean +/-standard deviation.

Glycerol viscosity. 5000X SYPRO Orange was diluted to 10 μM (“5X”) in each glycerol:ddH2O mixtures (mol/mol; Sigma-Aldrich G7893-2L) and mixed thoroughly. Then, 40 μL of each mixture was added to each well of a clear, flat-bottom 384-well plate (Greiner Bio-One ref 781091). Solutions were photographed in ambient daylight. On a Molecular Devices SpectraMax M5 fitted with a 384-well plate adaptor, absorbance spectra was measured from 350 – 700 nm. Emission following excitation at 485 nm was collected every 10 nm from 515 nm to 750 nm.

Structure and concentration of 5000X SYPRO Orange. From 200 μL of 5000X SYPRO Orange stock (Thermo Fisher Scientific Reference S6650, Lot 2008138), a dry red powder was produced by removal of DMSO under reduced pressure in the dark (GeneVac EZ-2 Elite, High BP setting, lamp off). Presented results are from six technical replicates over two separate experiments. LC-MS spectra were collected on an Agilent ACQ-TQD in 0.1% Formic Acid; NMR spectra were collected in d6-DMSO on a Bruker (400 MHz 1H/100 MHz ^13^C).

See Supplementary Methods for a detailed list of materials.

