## Supplemental Information for "Three Essential Resources to Improve Differential Scanning Fluorimetry (DSF) Experiments"

### Supplementary Information

#### Supplemental Tables


1 Properties of the proteins tested for the effects of heating rate. . 3
2 Parameters used to generate DSF data from Model 2 at varied heating rates. . . . . . . . . . . . . . . . . . . . . . . . . . . . . . . . 4
3 Diversified functions and commercial sources of the panel of compounds used for colloidal aggregation experiments. . . . . . . . . 4
4 Summary of starting estimates used in DSFworld model fitting. . 22
5 List of material sources and instruments used in the experiments. 23


#### Supplemental Figures


1 Model 2 reproduces heating-rate $Tm_a$ dependence. . . . . . . . . 5
2 Irreversible and reversible unfolding take place at the same temperatures for a diverse panel of proteins. . . . . . . . . . . . . . . 6
3 Empirical examples of failed SYPRO Orange detection. . . . . . 7
4 Anatomy of a plate compatibility test. . . . . . . . . . . . . . . . 8
5 Diversified chemical properties of the panel of compounds used for colloidal aggregation experiments. . . . . . . . . . . . . . . . . 9
6 Colloidal aggregation of small molecules in DSF. . . . . . . . . . 10
7 Influence of small molecules on protein-independent fluorescence correlates to colloidal aggregate abundance. . . . . . . . . . . . . 11
8 The structure of commercial SYPRO Orange. . . . . . . . . . . . 11
9 Characterization of commercial SYPRO Orange. . . . . . . . . . 12
10 Changes in visual pigmentation of 10 $\mu M$ ("5X") SYPRO Orange solutions with pH or small molecule colloidal aggregation. . . . . 13
11 Supplementary Figure 11. Performance of the four fits at DSFworld against the 347-curve test dataset. . . . . . . . . . . . . 14


#### Supplemental Notes


**1 Theoretical DSF model 15**
1.1 Part I: thermodynamic models of protein thermal unfolding. . . . 15
1.2 Part II: Incorporation of the influence of kinetics. . . . . . . . . 16
1.3 Part III: Incorporation of dye binding and fluorescent activation. 17

1.4 Considerations and caveats. . . . . . . . . . . . . . . . . . . . . . . 17

**2 DSFworld data analysis 18**

2.1 First derivative, single Tm . . . . . . . . . . . . . . . . . . . . . . . 19
2.2 Sigmoid fitting, four possible models. . . . . . . . . . . . . . . . . . 19
2.3 Considerations and caveats . . . . . . . . . . . . . . . . . . . . . . . 24

### Supplemental Tables

| Protein | MW (kDa) | Function | Published oligomeric state | ΔTma |
|---|---|---|---|---|
| Lysozyme | 14.3 | Lysosomal hydrolase | Monomer | 0.8 |
| Malate dehydrogenase (MDH) | 36.2 | Periportal/oxidative enzyme | Dimer | 3.1 |
| PPIF | 22.04 | Cyclophilin | Monomer | 3 |
| PPIE | 42 | Metabolic enzyme | Decamer | 4.5 |
| PerAB Early Doors Mutant | 38.632 | Circadian-regulator, clock-shift mutant | Dimer | 5.4 |
| Per2 | 38.632 | Circadian regulator | Dimer | 4.7 |
| Hsc70 | 70.898 | Molecular chaperone | Monomer | 2.2 |
| CHIP | 32.69 | E3 ligase | Dimer | 2.6 |

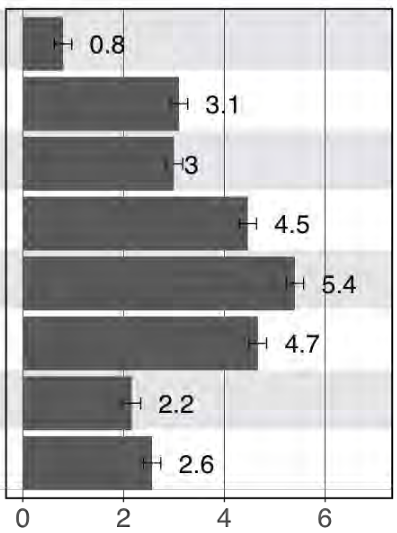


Supplemental Table 1: Properties of the proteins tested for the effects of heating rate.
Details on the representative protein panel from Figure 1, which were selected to sample different sizes, function, and oligomeric states. The range of $Tm_a$ values ($\Delta Tm_a$) determined at four heating rates ( 0.25, 0.5, 1 and 2 °C/min) is also shown. The error bars represent S.D. from at least two independent experiments.

| Parameter description | Parameter | Variable name in | Value | Varied or constant throughout Figure S2 |
|---|---|---|---|---|
| Heating rate | v | v_ | 0.5 - 2 ºC/min | Varied |
| Minimum temperature for significant irreversible unfolding | T* | T_star_ | 37 - 95 ºC | Varied |
| Enthalpy of reversible unfolding | ΔH | dHu_ | 200 kcal/mol | Constant |
| Thermodynamic melting temperature | $T_{1/2}$ | T_half_ | For panels a - c: 55 ºC<br>For panel d: 70 ºC | Constant |
| Heat capacity of reversible unfolding | $\Delta C_p$ | dCp_ | 8 kcal/mol | Constant |
| Activation energy of irreversible unfolding | $E_a$ | Ea_ | 150 kcal/mol | Constant |
| Dye detection of protein native state | $\text{Dye detection}_{\text{Native}}$ | nat_dye | 0 RFU/Mole fraction | Constant |
| Dye detection of protein reversibly unfolded state | $\text{Dye detection}_{\text{Rev. unfolded}}$ | unf_dye | 1 RFU/Mole fraction | Constant |
| Dye detection of protein irreversible unfolded state | $\text{Dye detection}_{\text{Irrev. unfolded}}$ | fin_dye | 1 RFU/Mole fraction | Constant |
| Temperature-dependent fluorescence decay | decay(T) | decay_rate | 0 RFU/ºC | Constant |
| Low temperature for integration in L(T) | $T_o$ | start_T | 25 ºC | Constant |
| High temperature for integration in L(T) | T | end_T | 25 - 95 ºC | Constant range |
| Final temperature for Temperature-dependent decay | $T_{final}$ | fin_T | 95 ºC | Constant |

Supplemental Table 2: Parameters used to generate DSF data from Model 2 at varied heating rates.
See Figure S2 for details.

| | Drug name | Citation | Typical use | Vendor | Catalog # | SMILES |
|---|---|---|---|---|---|---|
| 1 | Vemurafenib | 1 | anti-neoplastic melanoma | Cell Signaling Technology, Inc. | 17531S | ClC1=CC=C(C2=CN=C(NC=C3C(C4=C(F)C=CC(NS(CCC)(=O)=O)=C4F)=O)C3=C2)C=C1 |
| 2 | Miconazole | 1 | anti-fungal | Emolecules Inc | 501598605 | ClC1=CC=C(COC(C2=C(Cl)C=C(Cl)C=C2)CN3C=CN=C3)C(Cl)=C1 |
| 3 | Clotrimazole | 1 | anti-fungal | Emolecules Inc | 501544319 | ClC(C=CC=C1)=C1C(C2=CC=CC=C2)(N3C=NC=C3)C4=CC=CC=C4 |
| 4 | Ritonavir | 1 | anti-retroviral | Adipogen Corporation | 501687445 | O=C(NC(C(CC(NC(C(NC(N(CC1=CSC(C(C)C)=N1)C)=O)C(C)C)=O)CC2=CC=CC=C2)O)CC3=CC=CC=C3)OCC4=CN=CS4 |
| 5 | Emodin | 1 | natural product | Selleck Chemical Llc | 501362252 | OC1=C2C(C(C(C=C(O)C=C3O)=C3C2=O)=O)=CC(C)=C1 |
| 6 | Crizontinib | 2 | anti-non small cell lung carcinoma | Medchemexpress Llc | 501873893 | CC(C1=C(Cl)C=CC(F)=C1Cl)OC2=C(N)C=CC(C(C=N3)=CN3C4CCNCC4)=C2 |
| 7 | Sorafenib | 2 | advanced renal cell carcinoma | Medchemexpress Llc | 501871951 | CNC(C1=NC=CC(OC2=CC=C(NC(NC3=CC(C(F)(F)F)=C(Cl)C=C3)=O)C=C2)=C1)=O |
| 8 | Danazol | 1 | endometriosis | SIGMA-ALDRICH | D8399-100MG | CC1(C2)C(CCC3C1CCC4C3CCC4(O)C#C)=CC5=C2C=CO5 |
| | | | | | | ClC1=CC=C(C2=CN=C(NC=C3C(C4=C(F)C=CC(NS(CCC)(=O)=O)=C4F)=O)C3=C2)C=C1 |

Supplemental Table 3: Diversified functions and commercial sources of the panel of compounds used for colloidal aggregation experiments.
Citation 1: Colloidal aggregation: From screening nuisance to formulation nuance. 19, 188–200 (2018). [5]; Citation 2: Colloidal Aggregation Affects the Efficacy of Anticancer Drugs in Cell Culture. 7, 1429–1435 (2012) [10].

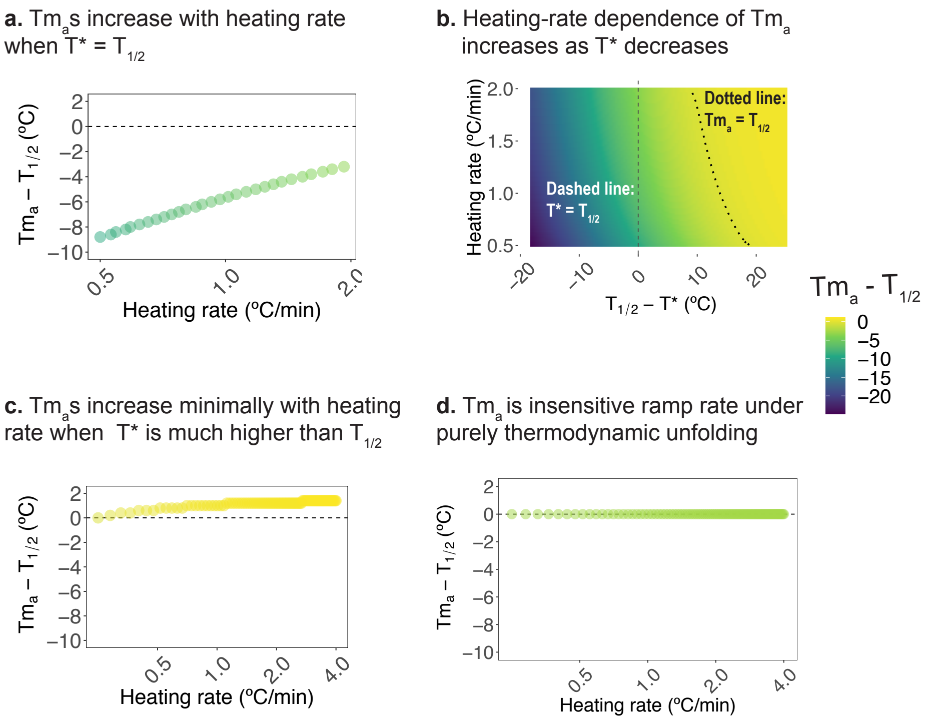


Supplemental Figure 1: Model 2 reproduces the dependence of $Tm_a$ on heating rate.

DSF data were computed from Model 2 for heating rates ranging from 0.5 - 2 °C/min and $T^*$ values ranging from 37 - 80 °C, keeping all other parameters constant (Supplementary Table 2). From the resulting data, $\text{Tm}_a$s were calculated using DSFworld by maximum of the first derivative (dRFU). **a.** When $\text{T}^* = \text{Tm}_{1/2}$, as suggested empirically by the results presented in Figure S2, $\text{Tm}_a$s increased by 6 °C as the heating rate was increased from 0.5 to 2 °C/minute, similar to the range observed empirically in Figure 1. **b.** The magnitude of the heating-rate dependent changes in $\text{Tm}_a$ increases as T* decreases. Additionally, when T* is sufficiently greater than $\text{Tm}_{1/2}$, $\text{Tm}_a$ can be coincidentally equal to $\text{Tm}_{1/2}$ at a particular heating rate, even in the presence of kinetic influences on unfolding (dotted line). **c.** When Model 2 parameters are adjusted to approximate lysozyme ($\text{Tm}_{1/2} = 70$ C, $\text{T}^* = 95$ C), the $\Delta$ $\text{Tm}_a$ associated with the change in ramp rate from 0.5 to 2 °C/min is 1 °C, similar to the 0.8 C observed in the experiment presented in Figure 1. **d.** In the case of thermodynamic unfolding, $\text{Tm}_a = \text{Tm}_{1/2}$, regardless of heating rate.

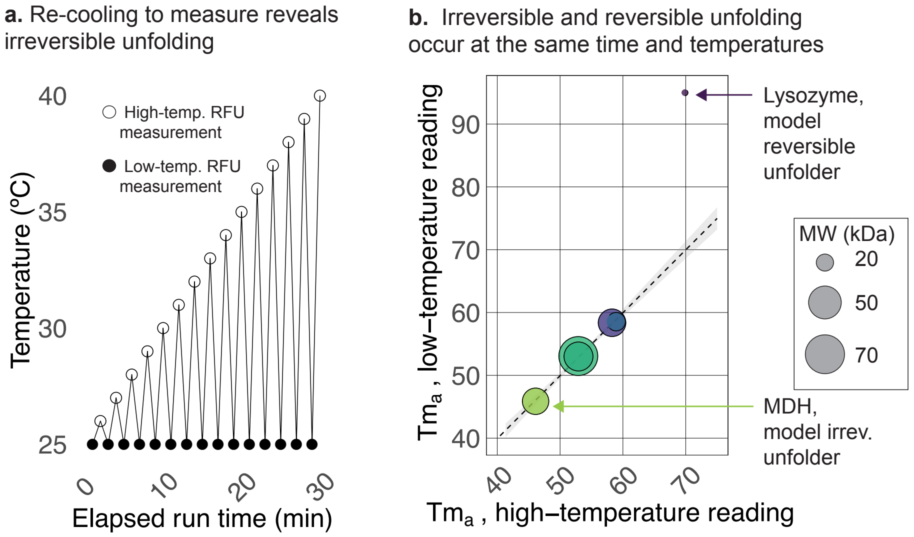


Supplemental Figure 2: Irreversible and reversible unfolding take place at the same temperatures for a diverse panel of proteins.
**a.** Schematic of the "up-down mode" thermocycling protocol used to emphasize the contribution of irreversible unfolding in a DSF experiment. Briefly, the irreversible state is maintained at low temperature, while the high temperature samples both reversible and irreversible unfolding. **b.** $Tm_a$ values produced from DSF experiments performed at either high-temperatures (reversible and irreversible) or low-temperatures (irreversible only) agree for all proteins except the model reversible-unfolding protein lysozyme, as expected. This result suggests that the temperature at which irreversible unfolding is significant (precisely, $k = 1/min$) is likely near or equal to the thermodynamic melting temperature, $T_{1/2}$.

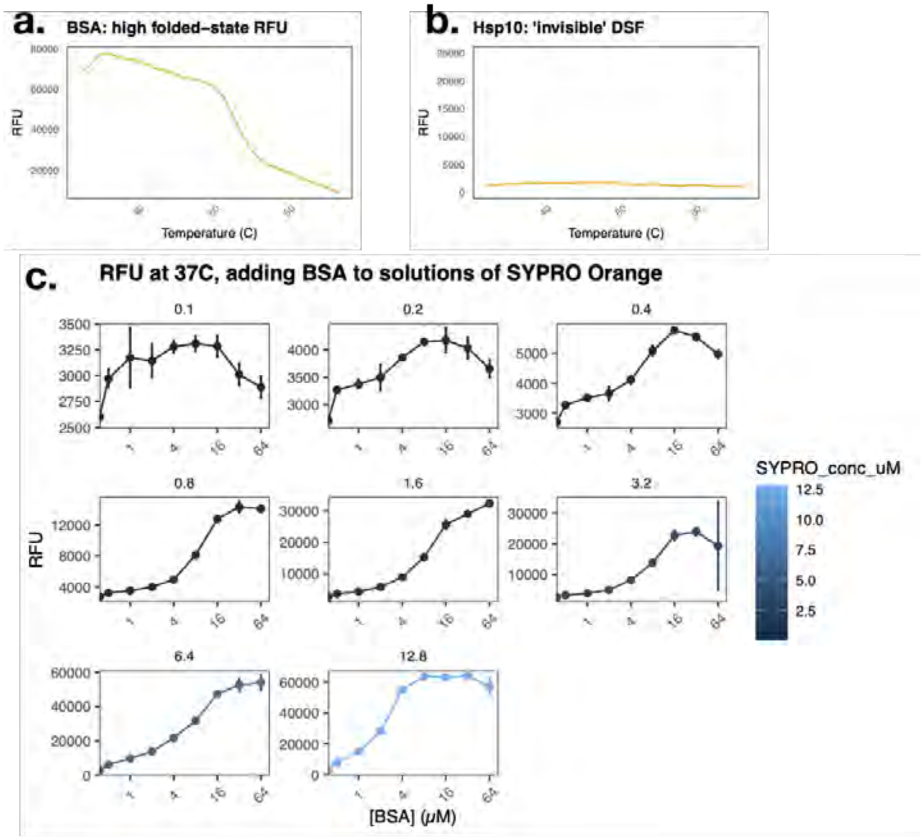


Supplemental Figure 3: Empirical examples of failed SYPRO Orange detection. **a.** SYPRO Orange detection of the folded state of Bovine Serum Albumin (BSA), likely due to SYPRO Orange binding to the native, folded state of the protein, leads to an obscured melting transition from which no $\mathrm{Tm}_a$ can be reliably obtained. **b.** SYPRO Orange does not detect unfolded 10 kDa heat-shock protein (Hsp10), likely due to no binding of SYPRO Orange to the unfolded state of the protein, leading to no discernible transition. Hsp10 is known to unfold within the measured temperature regime [3] **c.** SYPRO Orange fluorescence increases in a dose-dependent manner upon addition of native, folded BSA. The intensity of the induced SYPRO Orange fluorescence increases as with increasing concentrations of SYPRO Orange, from 0.1 to 12.8 $\mu M$. For determination of the concentration of SYPRO Orange, see Figure S8.

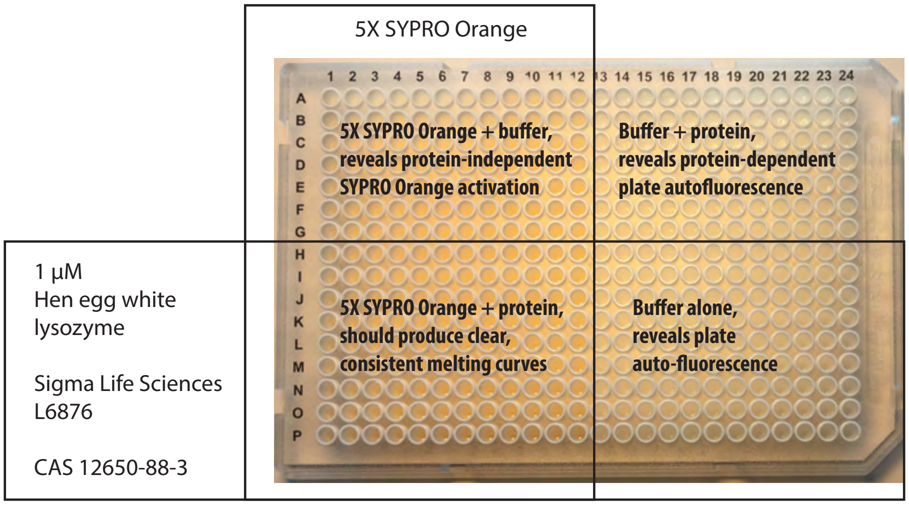


Supplemental Figure 4: Anatomy of a representative plate compatibility test. If a specific buffer will be used in downstream experiments, it is best to use that buffer for the plate compatibility test. Otherwise, any standard, simple buffer can be used, such as 10 mM HEPES, 200 mM NaCl, pH 7.20

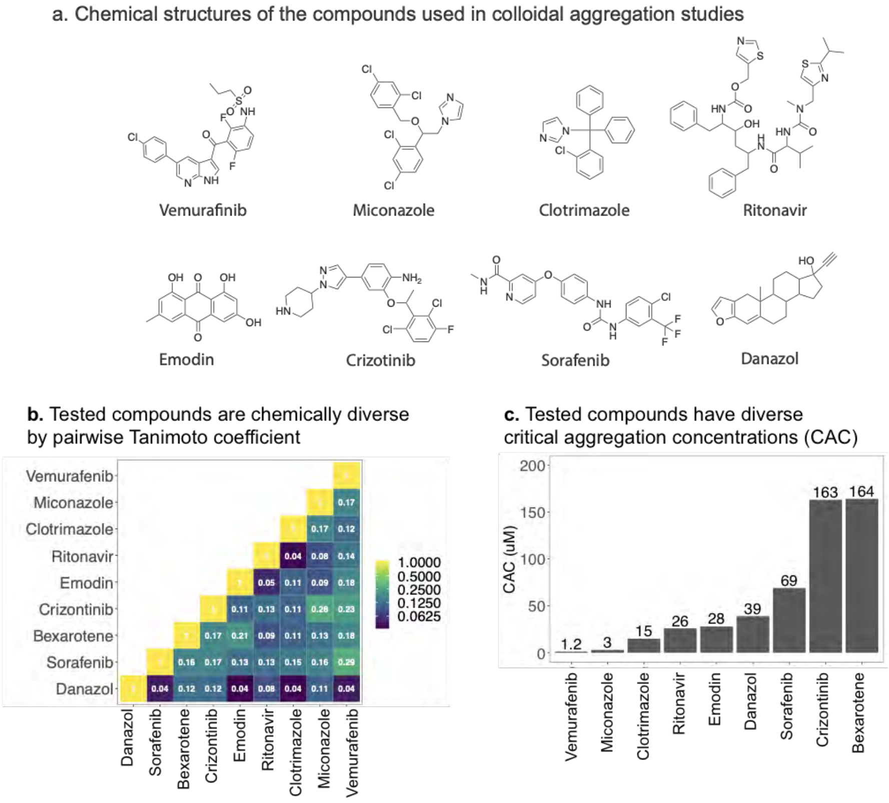


Supplemental Figure 5: Properties of the compounds used for colloidal aggregation experiments.
**a.** Chemical structures of the eight compounds in the panel. **b.** The compounds are structurally and chemically diverse, as described by low pair-wise AP tanimoto score coefficients, calculated by chemmineR package and online tools (https://chemminetools.ucr.edu/about/). **c.** The compounds cover a large range of critical aggregation concentration (CAC) values; values taken from references 1 and 2.

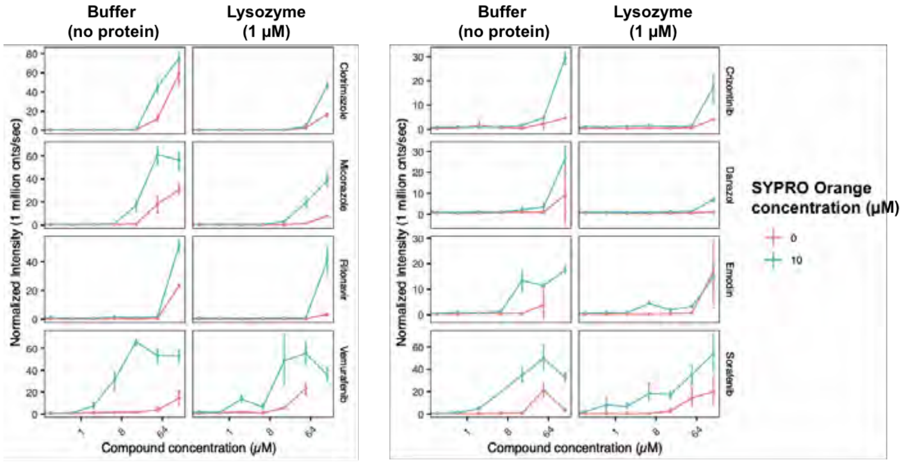


Supplemental Figure 6: Colloidal aggregation of small molecules as measured by DSF.

Dynamic Light Scattering (DLS) of the eight small molecules in the absence and presence of SYPRO Orange and 1 $\mu M$ lysozyme. All eight compounds (Clotrimazole, Miconazole, Vemurafenib, Crizotinib, Danazol, Emodin, Sorafenib) form colloidal aggregates within the tested concentrations (0 - 150 $\mu M$) for all conditions as determined by Dynamic Light Scattering (DLS) in buffer (10 mM HEPES, 200 mM NaCl, pH 7.20, 0.22 $\mu M$ filtered). In all cases, the presence of 5X SYPRO Orange ( 10 $\mu M$, teal lines) increased aggregation, while the addition of 1 $\mu M$ lysozyme reduced it.

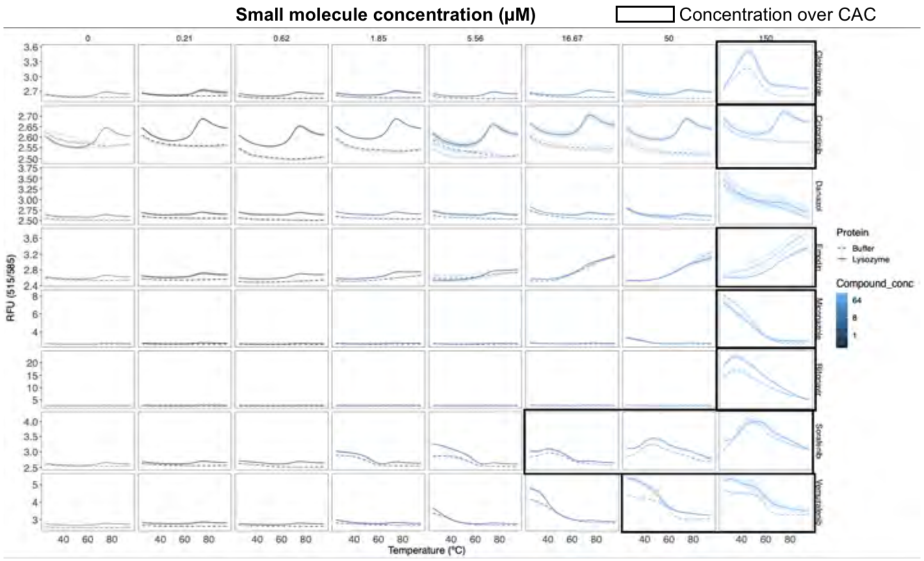


Supplemental Figure 7: Colloidal aggregates interfere with DSF signal, independent of protein.
At concentratinos above the CAC, aberrant dye fluorescence is observed by DSF in the presence of buffer, dye, and compound alone (dotted lines). The abberations are sufficient to obscure the melting signal of 1 $\mu M$ lysozyme (solid lines)..

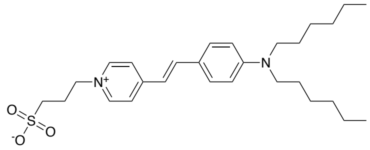


Molecular Weight: 486.72

Supplemental Figure 8: The structure of commercial SYPRO Orange.
LCMS (m/z) calculated 486.29, observed 487.0. This structure is consistent with that reported by Kroger et al 2017 [7]. Using the molecular weight of 486.29 g/mol for SYPRO Orange, the concentration of the 5000X stock was calculated to be 10.4 +/- 0.2 mM.

**a.**

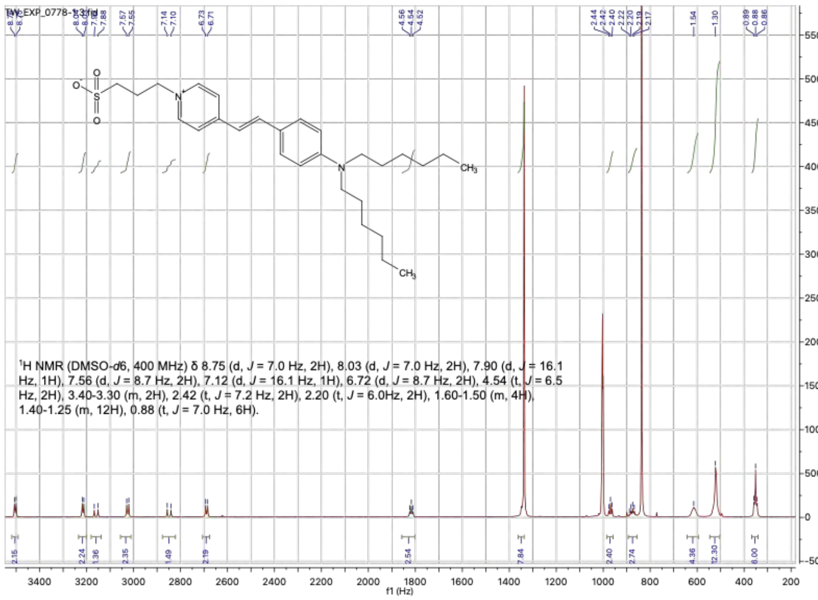


**b.**

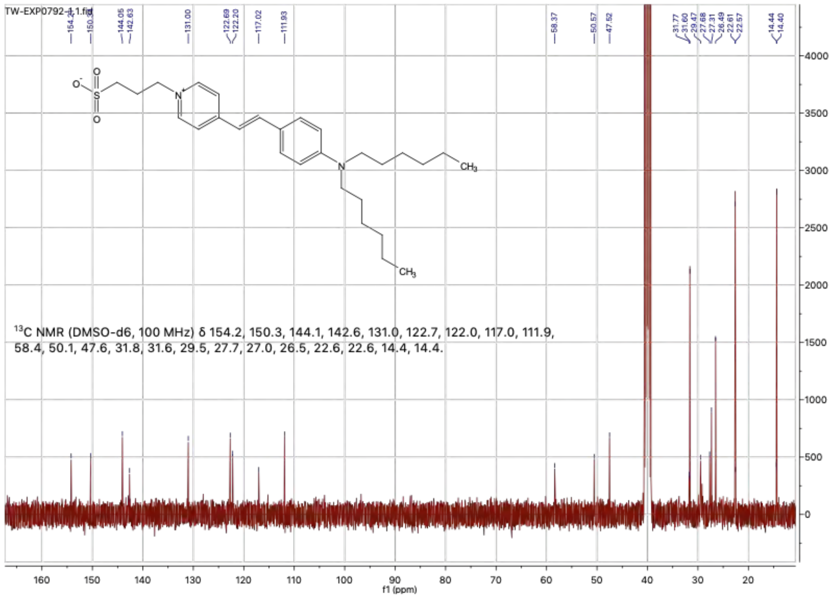


Supplemental Figure 9: Characterization of commercial SYPRO Orange.
The struture of commercial SYPRO Orange was determined by 1H-NMR and 13C-NMR. The molecular mass was confirmed by LCMS (m/z) calculated 486.29, observed 487.0. **a.** 1H-NMR of commercial SYPRO Orange. (DMSO-d6, 400 MHz) $\delta$ 8.75 (d, J = 7.0 Hz, 2H), 8.03 (d, J = 7.0 Hz, 2H), 7.90 (d, J = 16.1 Hz, 1H), 7.56 (d, J = 8.7 Hz, 2H), 7.12 (d, J = 16.1 Hz, 1H), 6.72 (d, J = 8.7 Hz, 2H), 4.54 (t, J = 6.5 Hz, 2H), 3.40-3.30 (m, 2H), 2.42 (t, J = 7.2 Hz, 2H), 2.20 (t, J = 6.0Hz, 2H), 1.60-1.50 (m, 4H), 1.40-1.25 (m, 12H), 0.88 (t, J = 7.0 Hz, 6H). **b.** 13C-NMR of commercial SYPRO Orange. (DMSO-d6, 100 MHz) $\delta$ 154.2, 150.3, 144.1, 142.6, 131.0, 122.7, 122.0, 117.0, 111.9, 58.4, 50.1, 47.6, 31.8, 31.6, 29.5, 27.7, 27.0, 26.5, 22.6, 22.6, 14.4, 14.4.

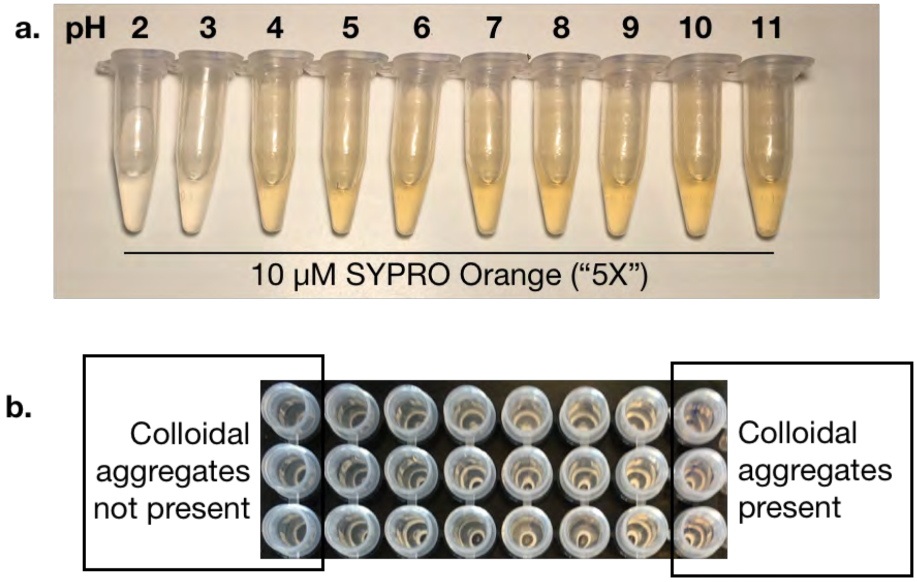


Supplemental Figure 10: Changes in visual pigmentation of 10 $\mu M$ ("5X") SYPRO Orange solutions with pH or small molecule colloidal aggregation.
a. 10 $\mu M$ solutions of SYPRO Orange, displaying decreased room temperature pigmentation in low-pH PBS. b. 10 $\mu M$ solutions of SYPRO Orange, displaying increased room temperature pigmentation in the presence of small molecule colloidal aggregates.

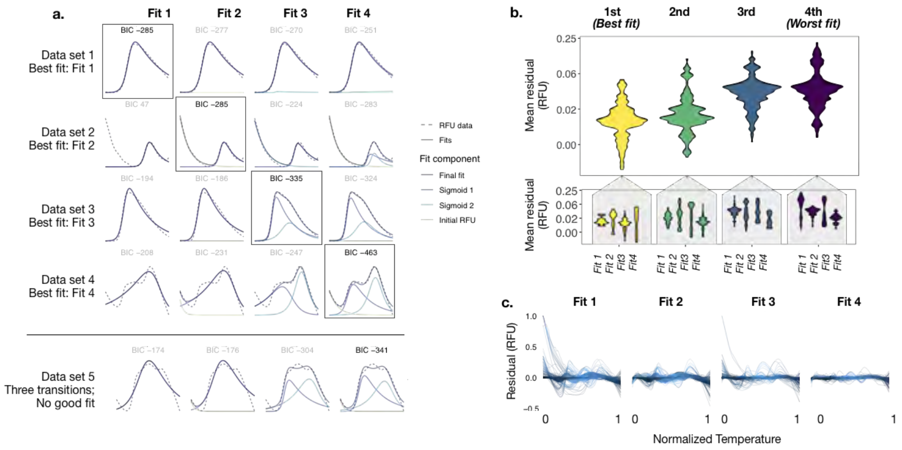

Supplemental Figure 11: Performance of the four fits at DSFworld against the 347-curve test dataset.

a. Representative curves best described by each of the four curve fitting options (fits 1-4). The best fit is defined as the one with the lowest Bayesian Information Criterion (BIC) for each data set, and outlined in black in the plot. Because DSFworld does not include a curve fitting option with three individual transitions, triple-transtion datasets like dataset 5 are not described by any of the four options. b. The distribution of mean residual RFUs for the 347-curve test data set, rank-ordered by BIC (1st is the best model for a given dataset, 4th is the worst), for any fit option (top panel), and broken out into each fit option (bottom panel). The decrease in mean residual with each additional model for this large dataset motivated the inclusion of the four fitting options available at DSFworld. c. Residual RFU for each of the 347 curves fit by each of the four options demonstrate that high residual values stem not from random noise, but the inability of an over-simplified fit option to describe a smooth feature in the data. For example, fits 1 and 3 cannot account for high initial fluorescence, so high initial residuals are regularly observed with these options; in these cases, fits 2 or 4 are often the best choice.

### 1 Theoretical DSF model

The purpose of the following note is to elaborate on Model 2, which while a useful framework for DSF in our experience, is not exhaustive. It is likely that individual DSF applications may require adjustments of improvements to this framework for maximum utility. To facilitate this, the following is provided as a starting point for these adjustments.

#### 1.1 Part I: thermodynamic models of protein thermal unfolding.

Typical DSF experiments assume a model where the protein has only two relevant states—native and unfolded—and these states are always at equilibrium [4, 8]. That is:

$$Native \leftrightharpoons Reversibly\ unfolded \qquad \text{(Model scheme 1)}$$

Through the 1970s and 1980s, the thermal unfolding of over 50 proteins was monitored by DSC [13, 2]. When considered in sum, these experiments suggested that the relationship between $\Delta G_{Fold}$ and temperature could be adequately described in terms of (1) the melting temperature of the protein $T_{1/2}$, (2) the enthalpy of unfolding ($\Delta H_{unfolding}$), and (3) the change in heat capacity $\Delta C_P$ associated with protein unfolding, using the empirically-derived relationship [13, 2, 14]:

$$\Delta G(T) = \Delta H_U \frac{T_{1/2} - T}{T_U} - \Delta C_p \left( T_{1/2} - T \left( 1 - ln \frac{T}{T_{1/2}} \right) \right) \qquad (1)$$

Equation 1 relates$\Delta G_{Fold}$ to temperature, which is converted to the fraction of protein folded ($X_{Folded}$) and unfolded ($X_{Unfolded}$) via the Gibbs-Hemholtz equation:

$$\Delta G = -RTln(K) \qquad (2)$$

re-arranged to:

$$K = e^{\frac{-\Delta G}{RT}} \qquad (3)$$

For two-state systems like Model scheme 1, the equilibrium constant K is related to the population of the native and reversibly unfolded states:

$$X_{Folded} = 1 - \frac{K}{K+1}$$

$$\text{(Model 1)}$$

$$X_{Reversibly\ unfolded} = \frac{K}{K+1}$$

Equation 1 is an established empirical relationship; equations 2 and 3 are general principles of thermodynamics. The thermodynamic model of unfolding built from them—"Model 1"—is displayed in the top panel of the interactive simulation in DSFworld. The outcomes of Model 1 respond only to changes in the thermodynamic properties of the protein–$\Delta H$ , the "true" (thermodynamic) Tm $T_{1/2}$ , and $\Delta C_P$–as expected for purely thermodynamic unfolding. $T_{1/2}$ controls the midpoint of the unfolding transition; $\Delta H$ and $\Delta C_P$ change the shape.

#### 1.2 Part II: Incorporation of the influence of kinetics.

The influence of kinetics is then included by adapting Model 1 to contain an irreversible unfolding step. That is:

$$Native \leftrightharpoons Reversibly\ unfolded \rightarrow Irreversibly\ unfolded \quad \text{(Model scheme 2)}$$

Model scheme 2 is the simplest of the classic Lumry-Eyring models of mixed thermodynamic-kinetic unfolding. Our incorporation of kinetic influence into thermodynamic unfolding models via the inclusion of a second irreversible unfolding step is adapted directly from previous work by Jose Sanchez-Ruiz extrapolating Lumry-Eyring models to DSC [15]. Briefly, the rate of conversion from the reversibly to irreversibly unfolded population is captured by a first-order rate constant $k$. The Arrhenius equation—a standard formula for the temperature-dependence of reaction rates—modifies $k$ as temperature increases in the simulation:

$$k\ (min^{-1}) = e^{-\frac{E_a}{R}\left(\frac{1}{T}-\frac{1}{T_{kin}}\right)} \tag{5}$$

Where $E_a$ is the activation energy of irreversible unfolding, $T$ the temperature in Kelvin, $T_{kin}$ a benchmark for the temperature at which irreversible unfolding becomes significant (precisely, where $k = 1 min^{-1}$ ), and R the gas constant 8.314 $\frac{J}{molK}$. The population of each state (e.g. folded, reversibly unfolded, irreversibly unfolded) is described by the two thermodynamic models from equation 3, joined now by an irreversibly unfolded state, each modified by the same temperature- and ramp rate-dependent kinetic parameter, $L(T)$:

$$X_{Folded} = 1 - \frac{K}{K+1}L(T)$$

$$X_{Rev.\ unf.} = \frac{K}{K+1}L(T) \tag{6}$$

$$X_{Irrev.\ unf.} = 1 - L(T)$$

Where $L(T)$ is obtained for each temperature $T$ by integration from a low temperature where k is negligible (in this work, $T_o = 298K$ ), followed by division by the thermocycling ramp rate in minutes, $\nu$, as below:

$$L(T) = 1 - e^{-M(T)} \tag{7}$$

where

$$M(T) = -\frac{1}{\nu}\int_{T_0}^{T} k\left(\frac{K}{K+1}\right) dT \tag{9}$$

As expected, in the absence of irreversible unfolding and therefore kinetic influence $k = 0$, which makes $L(T) = 1$ and $X_{Irrev.\ unf.} = 0$ for all temperatures, and equation 9 becomes equal to the thermodynamic models in Model 1.

#### 1.3 Part III: Incorporation of dye binding and fluorescent activation.

All of the modeling presented to this point applies broadly to heat-based denaturation of proteins, predicting the relative populations of the folded, reversibly unfolded, and irreversibly unfolded states of the protein. DSF data is simulated from these relative populations by multiplying the abundance of each state (native, reversibly unfolded, irreversibly unfolded) by the extent to which it activates the dye, comprising both dye binding and quantum yield:

$$RFU_{Folded} = X_{Folded} \times Dye\ detection_{Folded}$$

$$RFU_{Rev.\ unf.} = X_{Rev.\ unf.} \times Dye\ detection_{Rev.\ unf.} \tag{10}$$

$$RFU_{Irrev.\ unf.} = X_{Irrev.\ unf.} \times Dye\ detection_{Irrev.\ unf.}$$

The extent of dye activation resulting the three states is summed to produce the total signal:

$$RFU_{all} = (RFU_{Folded} + RFU_{Rev.\ unf.} + RFU_{Irrev.\ unf.}) \times decay(T) \tag{Model 2}$$

where

$$decay(T) = 1 - D \times \left(\frac{T}{T_{final}}\right) \tag{11}$$

The final modeled data, $RFU_{observed}$, is calculated from $RFU_{all}$ by multiplication by a linear temperature-dependent decay. This linear decay encompasses protein-independent loss of dye quantum yield and hydrophobic-driven binding at elevated temperatures.

#### 1.4 Considerations and caveats.

- There are many theoretical models of protein unfolding [15, 1, 16]. If the system of interest follows a different model of protein unfolding, then Model 2 will need to be adjusted. For example, Model 2 uses a first-order rate constant (equation 5) to describe the kinetically-controlled irreversible step, while aggregation-based irreversible unfolding processes, and particularly amyloid formation, are widely thought to follow higher-order kinetics [6].

- The $\Delta H_{Unfold}$ of the transition from the reversibly unfolded to the irreversibly unfolded state is assumed to be 0. This is currently a widespread assumption in the literature derived from DSC data [15, 11, 12, 9]. However, this is a caveat for the application of Model 2 to systems where $\Delta H_{Irrev.unf.}$ unfolding is known to be significant.

- Model 2 does not consider the influence of dye binding on the unfolding trajectory. In reality, dye-protein interactions must occur with some favorable $\Delta G_{binding}$, though both the magnitude and protein-to-protein consistency of this $\Delta G_{binding}$ is likely variable not clear at this time. The omission of dye binding energies from Model 2 should not affect outcomes for dye binding to the irreversibly unfolded state, as this transition is dominated by the associated activation energy (which is insensitive to the energy of the irreversibly unfolded state) rather than the relative $\Delta G$ between the reversibly and irreversibly unfolded states. However, dye binding to the reversibly unfolded state would perturb the $\Delta G_{unfolding}$, likely by stabilizing the reversibly unfolded state, thereby simultaneously decreasing the protein $Tm_a$ and increasing the activation energy of the irreversible, kinetically-controlled transition to the irreversibly unfolded state. Whether this dye-based stabilization of the reversibly unfolded state would lead to dye concentration-dependent increases or decreases in $Tm_a$ would likely depend on the extent to which the dye also detected the irreversibly unfolded state. Whether or not reversibly unfolded states are rendered irreversible in the context of DSF due to dye binding remains an interesting and unanswered question.

### 2 DSFworld data analysis

DSFworld analyzes raw DSF data. It accepts raw Temperature vs Fluorescence data, and exports visualizations and apparent melting temperatures. Most DSF data can be easily analyzed using either of two general approaches—first derivative or sigmoid fitting—and DSFworld supports both. These different analysis methods don't represent conflicting interpretations of DSF data; rather, they are different mathematical approaches which typically return the same $Tm_a$ value and can be used interchangeably.

The fitting methods may not work for some systems (see caveats below). To support customization and modification of the DSFworld analyses in these cases, the full code for the DSFworld website, as well as stand-alone R scripts, and modularized applications for data uploading, data formatting, plotting, $Tm_a$ determination, and downloading are available on Github.

The methods used to determine the apparent melting temperatures calculated on this website are as follows:

#### 2.1 First derivative, single Tm

Using this method, a single Tm is calculated for each DSF curve in the following manner:

1. The first derivative of the input data are calculated using a Savistky-Golay filter with a filter length of three degrees.
2. A Loess smoothing function is then used to interpolate the first derivative data to 0.1 C increments.
3. The maximum of the interpolated first derivative data is returned as the $\text{Tm}_a$.

The script and all associated functions to implement and modify this analysis outside of DSFworld is available on GitHub.

#### 2.2 Sigmoid fitting, four possible models.

To determine the number of models necessary to describe DSF data broadly, we first generated a representative dataset of DSF results. Briefly, we assembled a panel of X proteins diversified in molecular weight, biological activity, fold, and oligomeric state. We then performed DSF experiments in a variety of standard conditions, varying buffers, pH, concentrations of known ligands, SYPRO Orange concentrations, and heating rates. From these 347 DSF results, four archetypes of curves were visually identified: a single transition with no initial fluorescence (Model 1), a single transition with high initial, decaying fluorescence (Model 2), two transitions with no initial fluorescence (Model 3), and two transitions with high initial, decaying fluorescence (Model 4). These archetypes were mathematically defined as follows:

$$RFU(T) = Sig_1(T) \tag{Model 1}$$

$$RFU(T) = Sig_1(T) + Id(T) \tag{Model 2}$$

$$RFU(T) = Sig_1(T) + Sig_2(T) \tag{Model 3}$$

$$RFU(T) = Sig_1(T) + Sig_2(T) + Id(T) \tag{Model 4}$$

Where the general form of the decaying sigmoids, $Sig_i(T)$, is:

$$Sig_i(T) = \frac{A_i}{1 + e^{\frac{Tma_i - T}{scali}}} \times e^{d \times (T - Tma_i)}$$

- $Sig_i(T)$ is the RFU value at temperature $T$

- $A_i$ is the scaling factor for the final sigmoid
- $Tma_i$ is the $Tm_a$
- $scal_i$ controls the slope of the transition
- $d$ is the magnitude of the temperature-dependent RFU decay

And where the general form of the initial decaying fluorescence, $Id(T)$, is:

$$Id(T) = C \times e^{(T \times id)}$$

- $Id(T)$ is the RFU of the initial fluorescence at temperature $T$
- $C$ is the starting value of the initial fluorescence
- $id$ defines the rate and linearity of the decay from $C$

Approximately 10 datasets of each visual archetype were extracted and used throughout development of the curve fitting scripts. The resulting script was tested by fitting the 347-curve dataset, after which no further modifications were made to the script to maximize the relevance of the test results to the performance of the exact procedures applied at DSFworld. A stand-alone script for the model fitting is available at GitHub, alongside the 347-curve sample dataset used to test it.

The final fitting procedure is as follows:

1. A normalized version of the full uploaded dataset is generated by individually normalizing both raw RFU data and temperatures to a 0 to 1 range. This step minimizes inconsistent curve fitting behavior from arbitrary differences in measured temperature ranges and magnitudes of RFUs reported from different qPCR instruments.

2. From the normalized raw data, first- and second-order derivatives are calculated with respect to temperature using a Savitsky-Golay filter over a three-degree window (the number of individual measurements in a three-degree window is calculated from the non-normalized temperatures). A smoothed, interpolated version of the first- and second-derivative data are then calculated using a Loess filter of span 0.1 (10 percent of the measured temperature range).

3. Starting parameters for the models are generated in the following manner:

   - *$Tm_a$ of the major transition (All models)*
     In DSF, multiple transitions typically present as one high-magnitude (major) transition joined by a smaller (minor) one. In our experience, reasonable estimates for the $\mathrm{Tm}_a$ of the major transition can

typically be identified as the smooth major peak in the first derivative data; this is true for curves with both single and multiple transitions. Specifically, smooth peaks are identified in the Loess-smoothed first derivative data using the findPeaks() function from the quantmod package. Any peaks identified in the first or last five data points in the run are then removed because (i) in our experience, irreproducible, noise-like variations in DSF data are highest in these regions, meaning that such peaks are typically artifactual and (ii) reproducible transitions in DSF typically occur over at least a 10-measurement window, meaning that any transitions within the first or last five data points, if not artifactual, are likely partial. We have not yet encountered an application which required accurate fitting of partial curves in DSF (it is better to extend the measured temperature range, or optimize reaction conditions to bring the transition fully within the measured temperature range), so we did not optimize or test the performance of the DSFworld curve fitting in these cases. Finally, to eliminate peaks resulting from minor noise, any identified peaks with a maximum dRFU value of less than 0.0002 are removed as well. Peaks are then rank-ordered by their maximum dRFU, such that the largest peak in the dRFU is provided as the starting estimate for the $\mathrm{Tm}_a$ of the first sigmoid in subsequent model fitting.

- *$Tm_a$ of the minor transition (Models 3 and 4)*
  Estimation of the $\mathrm{Tm}_a$ of the minor transition are often more challenging. In DSF data, minor transitions often occur close in temperature to the major peak, appearing in the first derivative data as a shoulder on the primary peak. None of the peak finding algorithms we tested for use at DSFworld consistently identified these shoulders. However, we found that the minor transitions were more consistently captured as small stretches of positive-slope linearity in the raw data. These stretches can be quantified as valleys in the second derivative of the raw data. Specifically, smooth valleys are identified in the Loess-smoothed second derivative data using the findValleys() function from the quantmod. From the identified valleys, any which occur when the first derivative is $< 0$ are discarded. This step is necessary to separate the desired estimated $\mathrm{Tm}_a$s of the minor transitions from the largely linear, negatively-sloped post-maximum regions which occur in most DSF data. Any valleys which occur in the first or last five measurements are discarded for the same reasons described in the previous paragraph.

  To generate the final starting estimates for model fitting, the temperatures at which the identified peaks and valleys occur are combined. If multiple $\mathrm{Tm}_a$s are estimated within a three-degree window, only the lowest-temperature $\mathrm{Tm}_a$ of the closely-spaced estimates are retained. If only one $\mathrm{Tm}_a$ estimate is returned, a second $\mathrm{Tm}_a$ is estimated as the measurement directly after the first estimated $\mathrm{Tm}_a$.

- *Magnitude of initial, decaying fluorescence (Models 2 and 4)*
  The magnitude of the initial fluorescence is estimated as the first RFU value in the dataset.

- *Estimation of remaining parameters*
  The starting estimates for the remaining parameters are the same for all input data, because their empirical variation is, in our experience, relatively small. These are set as follows:

| Parameter | Parameter description | Starting estimate | Used in models... |
|---|---|---|---|
| Asym1 | Relative magnitude of major transition | 1.0 | 1, 2, 3, 4 |
| Asym2 | Relative magnitude factor for minor transition | 0.1 | 3, 4 |
| xmid1 | Tma of major transition | data-dependent | 1, 2, 3, 4 |
| xmid2 | Tma of minor transition | data-dependent | 3, 4 |
| scal | Slope of major transition | 0.03 | 1, 2, 3, 4 |
| scal2 | Slope of minot transition | 0.03 | 3, 4 |
| d | Temperature dependent RFU decay of major transition | -1 | 1, 2, 3, 4 |
| d2 | Temperature dependent RFU decay of minor transition | -2 | 3, 4 |
| id_d | Relative magnitude of initial fluorescence | data-dependent | 2, 4 |
| id_b | Temperature dependent RUF decay of initial fluorescence | -5 | 2, 4 |

Supplemental Table 4: Summary of starting estimates used in DSFworld model fitting.

4. From the resulting fits, the final parameters are then used to generate curves for each individual component of the full fit, multiplied by their individual temperature-dependent decay terms. These components are: the first sigmoid (all fits), the second sigmoid (fits 3 and 4), and high initial fluoresence (fits 2 and 4). The $Tm_a$ values are then calculated by taking the maximum of the first derivative of each isolated sigmoid component, as described section 2.1 of this supplementary note. This approach is analogous to the separation of a complex melting transition into its most likely individual unfolding populations, and then the use of the currently-accepted methods of $Tm_a$, which do not account for the influence of temperature-dependent decay on the midpoint of the transition, to each isolated population. This way, the DSFworld analysis leverages the temperature-dependent fluorescence decays necessary for robust fitting of complex transitions, while remaining consistent with the existing practices for DSF data analysis. The impact of correction for fluorescence decay on

| Product | Source | Reference | Lot |
|---|---|---|---|
| Hen Egg White Lysozyme, lyophilized powder | Sigma Life Sciences | L6876-10G | SLBT5161 |
| Bovine Serium Albumin, lyophilized powder | Sigma Life Sciences | Cat A2153-100G >96% | Lot # SLCB9433 |
| Glycerol | Sigma-Aldrich | G7893-2L | |
| DMSO | Sigma Aldrich | Prod 276855-100ml | Lot SHBK3913 |
| MicroAmp Optical Adhesive Film | Applied Biosystems | Ref 4311971 | Lot 201802294 |
| DSF plates (white 384-well micrometer) | Axygen | PCR-384-LC480WNFBC | Lot 09819000 |
| Incompatible DSF plates (white hard-shell 384-well micrometer) | BioRad | Cat HSR4805 | Control # 64072920 |
| 0.22 micron filter | Millex-GS | ref SLGS033ss | Lot # R8EA61590 |
| Glycerol experiment plates (black flat-bottom micro-amp) | Greiner Bio-one | ref 781091 | E170133P |
| DLS plates (low volume non-treated black, clear-bottom polystyrene) | Corning | ref 3540 | Lot # 00719006 |
| qPCR instrument | BioRad | CFX384 | |
| DLS instrument | Wyatt Technologies | DynaPro Plate Reader II | |
| Plate reader | Molecular Devices | SpectraMax M5 | |
| NMR | Bruker | Ascend 400 | |
| Genevac | Genevac | EZ-2 Elite | |
| LCMS instrument | Agilent Technologies | 1260 Infinity | |

Supplemental Table 5: List of material sources and instruments used in the experiments.

the model parameters themselves are discussed further in the Comments and Caveats section below.

5. Users of DSFworld can fit their data to any combination of the four models provided at DSFworld. The fit results of each model to all curves in the uploaded data can be downloaded individually. However, not all curves may be best described by the same model. By default, for summarized results in which only one fitted model per curve is desired, DSFworld will return the results from the fitted model with the lowest Bayesian Information Criterion (BIC) for each individual curve. The model of choice can also be manually selected by the user by clicking on the desired fit in the "select best model" plot in the analysis window.

#### 2.3 Considerations and caveats

- *Replicate handling.* Even when replicates of a particular condition have been defined by the user, a $\mathrm{Tm}_a$ is calculated for every individual curve, and the $\mathrm{Tm}_a$ for the condition is reported as the mean of the $\mathrm{Tm}_a$s calculated for each individual replicate, +/- standard deviation. When model fitting is used, if different models are selected for replicates of the same condition, results from the selected models with the smallest number of free parameters for that condition are applied to all members of that condition. Models 1, 2, 3, and 4 have four, six, eight, and ten free parameters, respectively.

- *Temperature-dependent fluorescent decay, and ramifications for the interpretation of model parameters.* DSFworld model fitting incorporates a temperature-dependent fluorescence decay coefficient throughout the entire the curve. At this time, we believe this approach is appropriate for the following reasons:

  - Fluorescence decay is not observed in low-temperature readings in up-down mode experiments. In an assembled panel of seven diverse proteins, fluorescence was monitored at both the high and low temperatures of an up-down mode experiment (Figure S2), and the results were compared to the results from straight-ramp experiments presented in Figure 1. While the $\mathrm{Tm}_a$s determined for the high- and low-temperature readings were similar, fluorescence decay was observed only in the high-temperature readings, and not in the low-temperature readings. Furthermore, the observed decay in the high-temperature readings of the up-down mode experiments matched that of the straight-ramp experiments. Combined, this observation suggests that the observed fluorescence decay in these experiments may be largely temperature dependent.
  - The stereotypical linearity of the post-maximum decay in real DSF data conflicts with a model in which this decay is caused by the transition of the protein into a final aggregated state which excludes

the dye. While this dye-exclusion model may certainly be true for some proteins, given this consistent linearly, we do not think it is an appropriate assumption for general-purpose DSF fitting tools.

- The strength of the protein-small molecule interactions is often decreased with temperature. Therefore, if the current hypothesis that SYPRO-protein interactions are primarily hydrophobic in nature is correct, an associated temperature dependent loss of dye binding is expected, even if all dye binding sites are retained as temperature is increased. This is anecdotally supported by observations that when SYPRO fluorescence is induced by the folded state of a protein, this fluorescence exhibits a very similar decay during heating through temperatures at which the protein undergoes no conformational changes–and by extension, changes in dye binding sites–as measured by circular dichroism (unpublished data).

The application of a temperature-dependent decay to the full sigmoid creates the following behaviors, of which users interested in direct interpretation of the parameters returned from DSFworld fits (as opposed to the $Tm_a$s returned from fits, which are robust to these effects)should be aware:

- $\mathrm{Tm}_a$s determined by sigmoid fits which incorporate temperature-dependence fluorescence decay will be higher than those calculated by methods which do not, such as the maximum of the first derivative. For the majority of DSF data, this difference is 1.5 C. Steeper decays produce larger differences.
- Two DSF curves may have the same $\mathrm{Tm}_a$ when analyzed using the maximum of the first derivative. However, if the two curves have different decay intensities, methods which incorporate temperature-dependent fluorescence decay, such as the sigmoid fits provided at DSFworld, will report different $\mathrm{Tm}_a$s for the two curves. This difference is typically small ($< 2$ C ), but increases as the difference in decay intensities increases.

It is possible that the above effects manifest in real DSF data, and are systematically overlooked by current analysis methods. If so, the correction of temperature-dependent decays in standard $Tm_a$ calculations may be appropriate. However, more information on the mechanisms and consistent behaviors of temperature-dependent decays in DSF will be necessary to answer this question broadly.
